# The *Vibrio cholerae* cytotoxin MakA induces noncanonical autophagy resulting in the spatial inhibition of canonical autophagy

**DOI:** 10.1101/2020.07.07.191411

**Authors:** Dale P. Corkery, Aftab Nadeem, Kyaw Min Aung, Ahmed Hassan, Tao Liu, Ramón Cervantes-Rivera, Alf Håkon Lystad, Hui Wang, Karina Persson, Andrea Puhar, Anne Simonsen, Bernt Eric Uhlin, Sun Nyunt Wai, Yao-Wen Wu

## Abstract

Autophagy plays an essential role in the defence against many microbial pathogens as a regulator of both innate and adaptive immunity. Among some pathogens, sophisticated mechanisms have evolved that promote their ability to evade or subvert host autophagy. Here, we describe a novel mechanism of autophagy subversion mediated by the recently discovered *Vibrio cholerae* cytotoxin, MakA. pH-dependent endocytosis of MakA by host cells resulted in the formation of a cholesterol-rich endolysosomal membrane aggregate in the perinuclear region. Aggregate formation induced the noncanonical autophagy pathway driving unconventional LC3 lipidation on endolysosomal membranes. Subsequent sequestration of the ATG12-ATG5-ATG16L1 E3-like enzyme complex required for LC3 lipidation at the membranous aggregate resulted in an inhibition of both canonical autophagy and autophagy-related processes including the unconventional secretion of IL-1β. These findings identify a novel mechanism of host autophagy subversion and immune modulation employed by *V. cholerae* during bacterial infection.

## Introduction

Macroautophagy (hereafter referred to as autophagy) is a highly conserved intracellular degradation system through which cytoplasmic constituents are sequestered in double-membraned autophagosomes and delivered to lysosomes for clearance. In addition to maintaining cellular homeostasis, autophagy plays a prominent role in the cellular response to microbial pathogens as a mediator of both innate and adaptive immunity (Deretic et al., 2013). From the direct sequestration and elimination of intracellular pathogens (xenophagy) (Levine, 2005), to the regulation of antigen presentation (English et al., 2009; Schmid et al., 2007) and cytokine secretion (Dupont et al., 2011; Zhang et al., 2015), the known roles for autophagy in the detection and elimination of cellular pathogens are numerous. As a result, many pathogens have evolved sophisticated mechanisms to evade or subvert host autophagy (McEwan, 2017; Wu and Li, 2019).

*Vibrio cholerae*, the causative agent of the acute diarrheal disease cholera, is a highly motile Gram-negative bacterium existing naturally in aquatic environments (Clemens et al., 2017). As an extracellular pathogen, its ability to manipulate host cell function depends on the secretion of numerous exotoxins capable of entering host cells and altering cellular processes in favor of bacterial infection. Recently, we described a novel cytotoxin of *V. cholerae*, the motility associated killing factor A (MakA), which we found to act as a potent virulence factor contributing to pathogenicity in the model host organisms zebrafish and *Caenorhabditis elegans* (Dongre et al., 2018). The *makA* gene was observed in all *V. cholerae* strains including non-O1 non-O139 *V. cholerae* (NOVC) isolates. Most NOVC strains lack the cholera toxin gene (*ctxAB*) and do not cause the typical diarrheal disease, cholera, but have recently been emerging as extra-intestinal pathogens (Faruque et al., 1998). Here we characterize the molecular basis of the interaction between MakA and mammalian host cells. We found that MakA binding to the plasma membrane of host cells promoted endocytosis resulting in the formation of an endolysosomal membrane-rich aggregate at the perinuclear region of intoxicated cells. The aggregate was targeted by the noncanonical autophagy pathway mediating LC3 lipidation on perturbed endosomal membranes. The end result is a spatial inhibition of canonical autophagy pathway by sequestering core autophagic components to the membranous aggregate. The spatial inhibition of autophagy was shown to reduce the unconventional secretion of IL1-β suggesting a novel approach to immune modulation by a bacterial toxin.

## Results

### MakA alters the cellular cholesterol distribution

MakA was recently identified in *V. cholerae* as a novel cytotoxin affecting both vertebrate and invertebrate hosts (Dongre et al., 2018), yet the molecular basis for its role as a virulence factor remained unknown. To determine the effect MakA has on human host cells, human colon carcinoma cells (HCT8) were treated with purified MakA at a dose approximately equal to the established IC50 (Figure S1) and RNA sequencing analysis was performed to monitor gene expression in a comparison with untreated cells (Figure 1A). Gene ontology (GO) enrichment analysis of differentially expressed genes identified several biological processes related to lipid metabolism and homeostasis with the ‘cholesterol biosynthetic process’ being the most significantly modulated (Figure 1B, C). Maintaining cholesterol homeostasis is essential to sustain cellular function. As such, elaborate regulatory networks exist to sense and maintain cellular cholesterol levels within a narrow range (Luo et al., 2020). To determine whether the observed transcriptional response was related to altered cellular cholesterol levels, HCT8 cells were treated with toxin and total cellular cholesterol was quantified. While MakA treatment did result in a slight dose-dependent increase in total cellular cholesterol (Figure 1D), visualization of free cholesterol revealed a striking toxin-induced redistribution, with large cholesterol aggregates visible at the perinuclear region of intoxicated cells (Figure 1E). A similar cholesterol redistribution was observed in cells treated with supernatant harvested from cultured *V. cholerae* (Figure S2).

**Figure 1.**
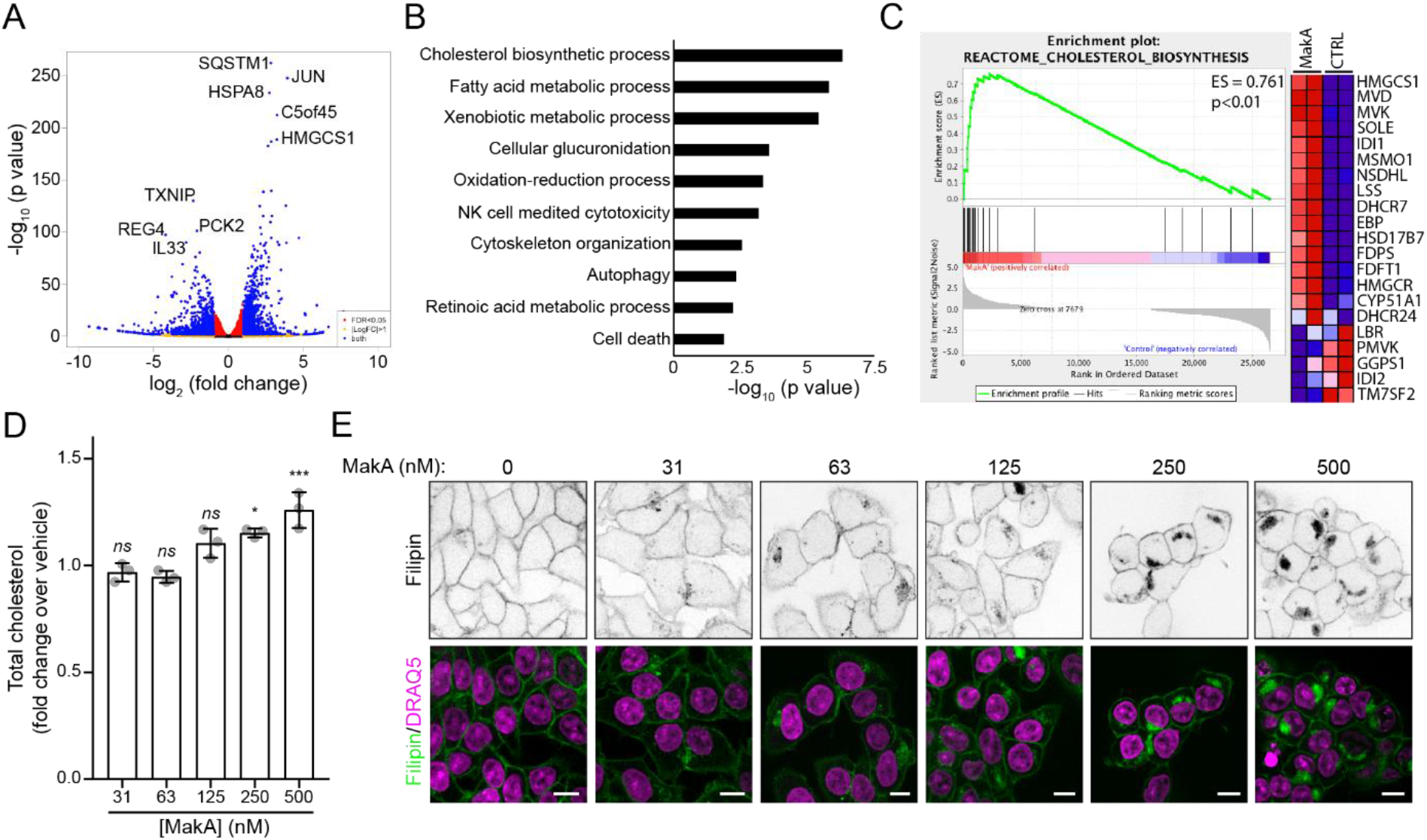
MakA treatment alters cellular cholesterol distribution. (**A**) Volcano-plot showing differentially expressed genes in HCT8 cells treated with 500 nM MakA for 48 h. Most significantly up and down regulated genes are highlighted. (**B**) Top ten gene ontology (GO) terms from GO enrichment analysis using differentially expressed genes from A. (**C**) Gene Set Enrichment Analysis (GSEA) of genes implicated in cholesterol biosynthesis. (**D**) Quantification of total cellular cholesterol (Amplex Red) in HCT8 cells treated with the indicated concentration of MakA for 24 h. Data points represent three biologically independent experiments; bar graphs show mean ± s.d. Significance was determined from biological replicates using a one-way analysis of variance (ANOVA) with Dunnett’s post-test against vehicle control. *p=0.0102, ***p=0.0002, ns=not significant. (**E**) Intracellular free cholesterol localization following 24 h MakA treatment as detected by Filipin staining. Nuclei were counterstained with DRAQ5. Scale bars, 10 μm.

### MakA-induced endocytosis promotes the formation of an endolysosomal membrane-rich aggregate

To determine the composition of the MakA-induced cholesterol-rich aggregates, and gain insight into the subcellular localization of MakA, correlative light and electron microscopy (CLEM) was performed on cells treated with Alexa Fluor 568(A568)-labeled MakA (Figure 2A). The toxin was found to accumulate at both the plasma membrane and within the perinuclear aggregate, which appeared membrane rich and contained vesicular structures closely resembling endosomes (Klumperman and Raposo, 2014). Immunofluorescence analysis of MakA treated cells using antibodies against the early, late, and recycling endosomal markers EEA1 (Simonsen et al., 1998) (Figure 2B, Figure S3), Rab7 (Feng et al., 1995) (Figure S3) and Rab11 (Urbé et al., 1993) (Figure S3) confirmed endosomal membrane-enrichment within the membranous aggregate. Aggregates were also shown to contain plasma membrane-localized transmembrane proteins Caveolin-1 (Cav1) (Figure S4A) and Na/K-ATPase α1 (Figure S4B), but were free of intracellular golgi (Figure S4C) or mitochondrial (Figure S4D) membrane. Furthermore, live-cell imaging of Cos7 cells transiently-transfected with the early endosomal marker EGFP-Rab5 (Bucci et al., 1992) and treated with A568-MakA found the toxin to be present within Rab5-positive endosomes (Figure 2C and Supplementary Video 1) suggesting MakA travels from the plasma membrane to the perinuclear membranous aggregate via endocytosis. To confirm endosomal involvement, Cos7 cells were transfected with a dominant-negative Rab5-S34N mutant (Li and Stahl, 1993). Expression of the mutant allele was sufficient to block A568-MakA-induced aggregate formation (Figure 2D).

**Figure 2.**
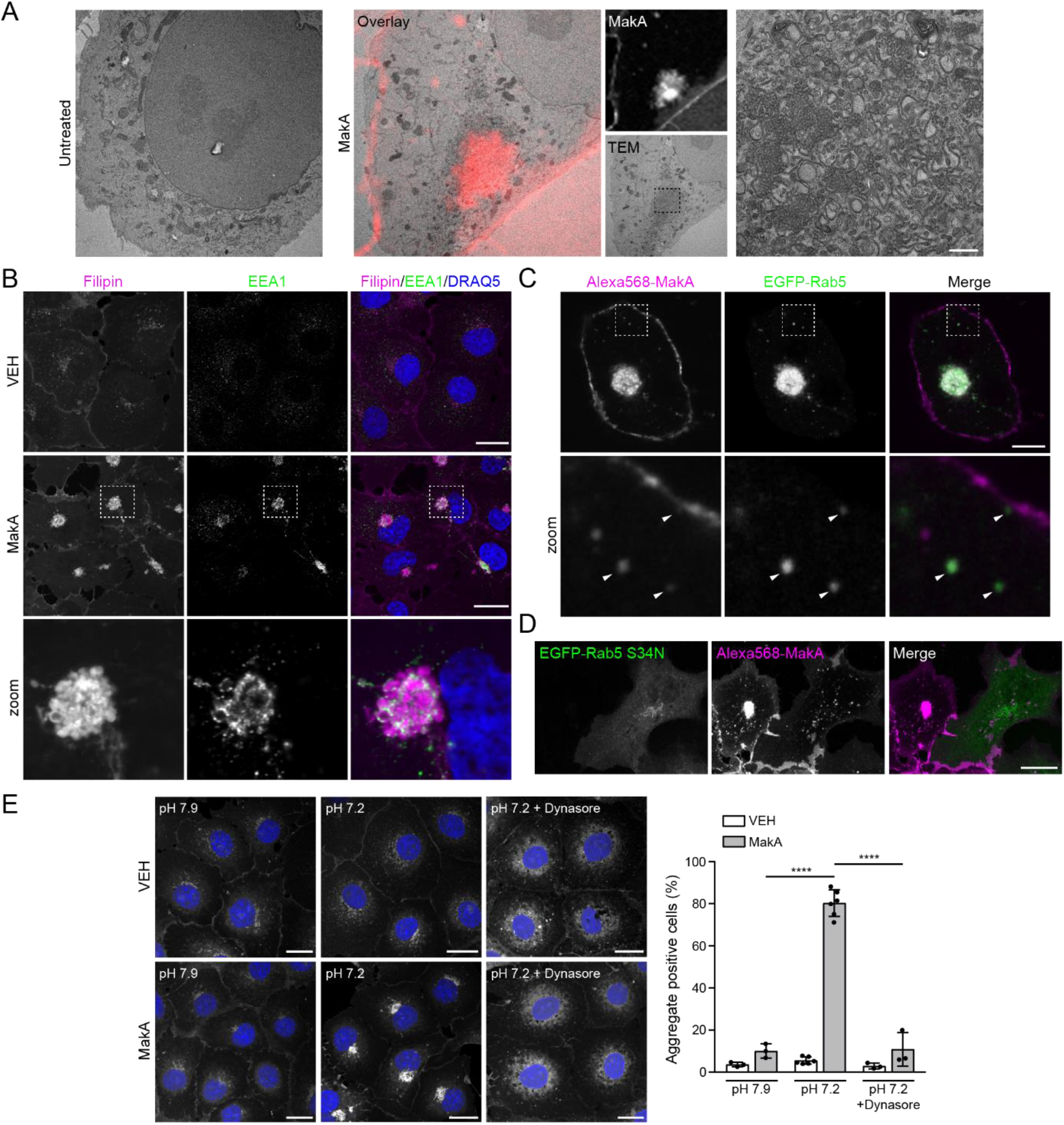
MakA-induced aggregates are formed as a result of pH-dependent endocytosis. (**A**) Representative correlative light and electron micrographs of untreated and A568-MakA (48 h, 500 nM) treated Cos7 cells. Scale bar, 500 nm. (**B**) Cos7 cells treated with vehicle (VEH) or 250 nM MakA for 48 h. Cells were stained with filipin to visualize MakA-induced aggregates and immunofluorescence performed with an antibody against endosomal marker EEA1. Nuclei were counterstained with DRAQ5. Scale bars, 20 μm. (**C**) Cos7 cells transfected with EGFP-Rab5 and treated with 250 nM A568-MakA for 48 h. Arrowheads indicate endosomes stained positive for Rab5 and MakA. Scale bar, 10 μm. (**D**) Cos7 cells transfected with EGFP-Rab5 S34N and treated with 250 nM A568-MakA for 48 h. Scale bar, 20 μm. (**E**) (left) Cos7 cells treated with 250 nM MakA for 24 h in media adjusted to the indicated pH with or without 80 μM dynasore. Nuclei were counterstained with DRAQ5. Scale bars, 20 μm. (right) Quantification of the percentage of cells containing aggregates. Data points represent biologically independent experiments (n > 100 cells per experiment); bar graphs show mean ± s.d. Significance was determined from biological replicates using a two-way ANOVA with Tukey’s multiple comparisons tests. ****p<0.0001.

Interestingly, MakA-induced endocytosis was found to be pH-dependent. Unlike HCT8 cells which show robust aggregate formation after 24 h of MakA treatment (Figure 1E), Cos7 cells failed to develop MakA-induced aggregates until 48 h post toxin treatment. This response time could be decreased by adjusting the starting pH of the culture media from pH 7.9 to pH 7.2 (Figure 2E). This pH-dependent aggregate formation could be prevented by treating cells with the dynamin inhibitor dynasore (Macia et al., 2006) to block dynamin-dependent endocytosis (Figure 2E). Together, these data confirm that the MakA-induced aggregate is formed as a result of toxin-induced dynamin- and Rab5-dependent endocytosis and is composed of plasma membrane-derived, cholesterol-rich endosomal membrane.

### MakA-induced aggregates are targeted by autophagy

RNA sequencing identified a significant change in expression of genes involved in the regulation of autophagy after toxin treatment (Figure 1B). To determine how MakA affects cellular autophagy, HCT8, HEK293T and Cos7 cells were treated with toxin and LC3 lipidation status was evaluated using western blot analysis (Klionsky et al., 2016). Upon the induction of autophagy, cytosolic LC3 (LC3-I) is conjugated to phosphatidylethanolamine (PE) to form membrane bound LC3-II (Tanida et al., 2008). A strong dose- and time-dependent increase in LC3 lipidation was observed following MakA treatment in all cell lines tested (Figure 3A, B). As was reported for MakA-induced endocytosis (Figure 2E), the timing of LC3 lipidation could be controlled by altering the starting pH of the culture media (Figure 3C). Interestingly, MakA-induced LC3 lipidation was restricted to within a narrow pH window (7.5 - 6.6). Dropping below pH 6.6 prevented LC3 lipidation and inhibited cholesterol-rich aggregate formation (Figure 3D).

**Figure 3.**
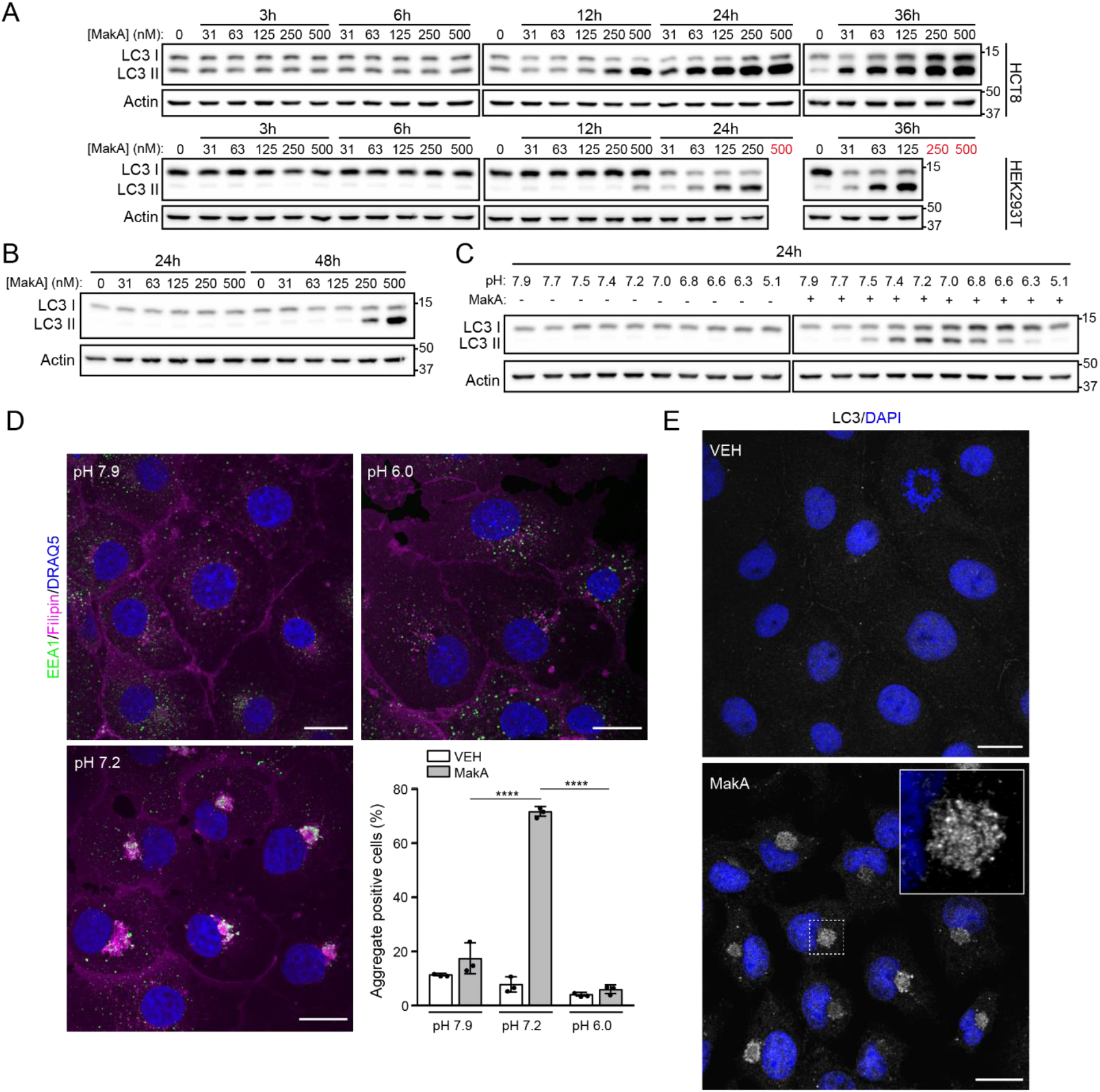
MakA treatment induces autophagy. (**A**, **B**) Western blot analysis of LC3 lipidation in HCT8, HEK293T and Cos7 cells treated with MakA, as indicated. Red values indicate toxic doses. Data are representative of at least three independent experiments. (**C**) Western blot analysis of LC3 lipidation in Cos7 cells treated with 250 nM MakA for 24 h in media adjusted to the indicated pH at the time of treatment. (**D**) Cos7 cells treated with 250 nM MakA for 24 h in media adjusted to the indicated pH at the time of treatment. Cells were stained with filipin to visualize MakA-induced aggregates and immunofluorescence performed with an antibody against endosomal marker EEA1. Nuclei were counterstained with DRAQ5. Scale bars, 20 μm. (bottom right) Quantification of the percentage of cells containing fillipin aggregates. Data points represent biologically independent experiments (n > 150 cells per experiment); bar graphs show mean ± s.d. Significance was determined from biological replicates using a two-way ANOVA with Sidak’s multiple comparisons tests. ****p<0.0001. (**E**) Cos7 cells treated with vehicle or 250 nM MakA for 24 h in pH 7.2 media. Immunofluorescence was performed with an antibody against LC3B. Nuclei were counterstained with DAPI. Scale bars, 20 μm.

To explore the relationship between aggregate formation and the induction of autophagy, we performed immunofluorescence analysis for endogenous LC3 in Cos7 cells treated with MakA. The majority of LC3 was found to be localized at the perinuclear aggregate after MakA treatment (Figure 3E). The aggregate was also enriched in mono- and poly-ubiquitinylated conjugates (Figure 4A); a common prerequisite for substrate recognition in selective autophagy (Kim et al., 2008; Pankiv et al., 2007), and the LC3-ubiquitin adaptor protein p62 (Figure 4B). Interestingly, both ubiquitinylated conjugates and p62 localized primarily to the periphery of the aggregate indicating that the aggregate could be targeted by the autophagic machinery after formation. We also observed clustering of lysosomes around the aggregate (Figure 4C), similar to what has been described for autophagy-mediated degradation of abnormal polypeptide aggregates (aggresomes) (Zaarur et al., 2014).

**Figure 4.**
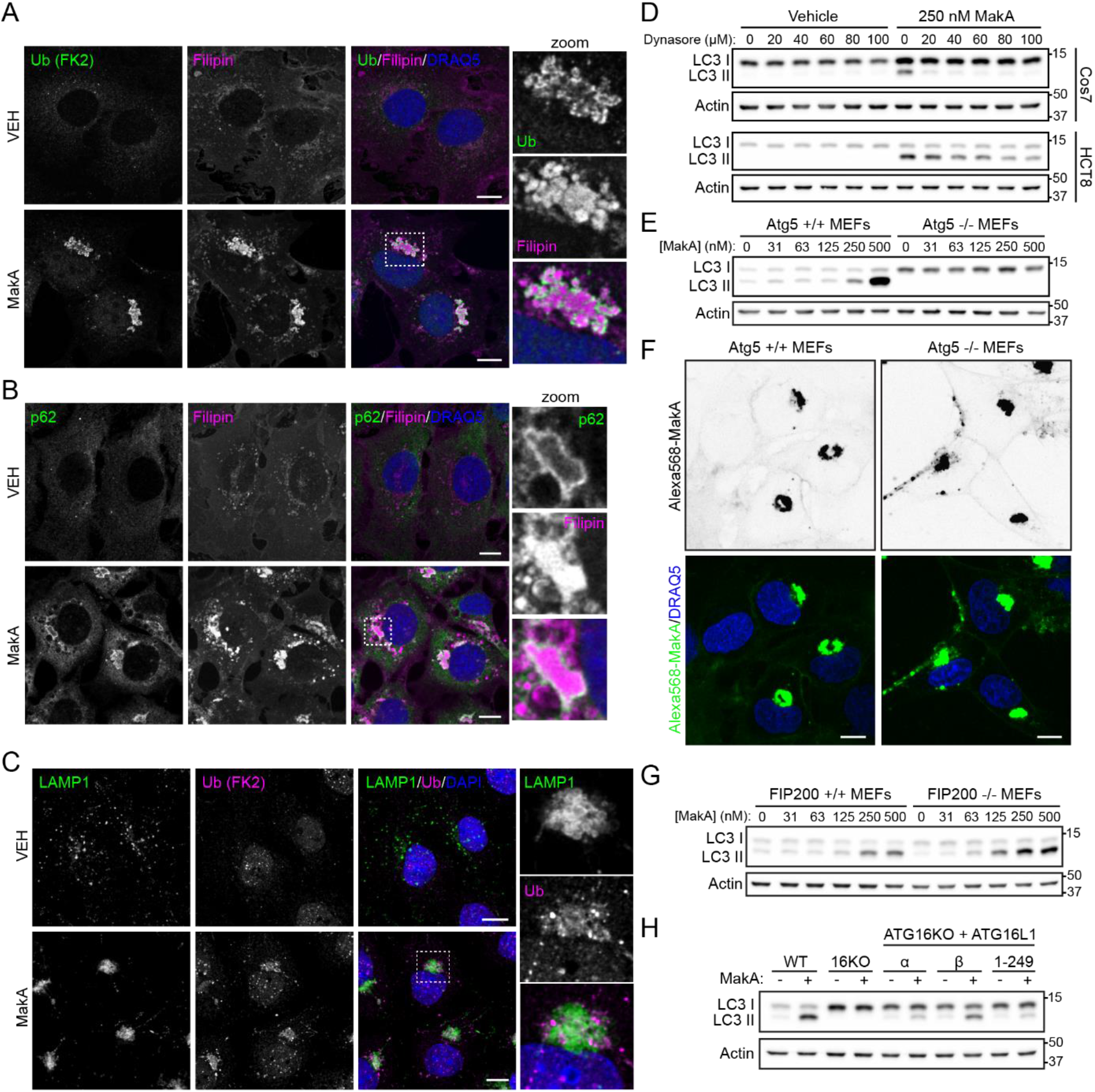
MakA-induced aggregates are targeted by the autophagic machinery. (**A**, **B**) MEFs treated with vehicle or 500 nM MakA for 24 h. Cells were stained with filipin to visualize MakA-induced aggregates and immunofluorescence performed with antibodies against mono- and polyubiquitinylated conjugates (A) or p62 (B). Nuclei were counterstained with DRAQ5. Scale bars, 10 μm. (**C**) Cos7 cells treated with vehicle or 250 nM MakA for 48 h. Immunofluorescence was performed with antibodies against mono- and polyubiquitinylated conjugates (Ub (FK2)) to identify aggregates, and LAMP1 to visualize lysosomes. Nuclei were counterstained with DAPI. Scale bars, 10 μm. (**D**) Western blot analysis of Cos7 and HCT8 cells treated with 250 nM MakA (pH 7.2) in the presence of increasing concentrations of dynasore. Data are representative of two independent experiments. (**E**) Western blot analysis of WT (Atg5+/+) and autophagy deficient (Atg5−/−) MEFs treated with the indicated concentration of MakA for 24 h. Data are representative of three independent experiments. (**F**) WT and autophagy deficient MEFs treated with 500 nM Alexa568-MakA for 24 h. Nuclei were counterstained with DRAQ5. Scale bars, 10 μm. (**G**) Western blot analysis of FIP200+/+ and FIP200−/− MEFs treated with the indicated concentration of MakA for 24 h. Data are representative of two independent experiments. (**H**) Western blot analysis of LC3 lipidation status in WT and ATG16L1-KO HEK293 cells rescued or not with ATG16L1α, ATG16L1β or ATG16L1 1-249 and treated with or without 250 nM MakA for 24 h. Data are representative of three independent experiments.

To confirm that autophagy was induced in response to aggregate formation, dynamin-dependent MakA endocytosis was inhibited with dynasore to prevent aggregate assembly (Figure 2E). Blocking aggregate formation was sufficient to prevent MakA-induced LC3 lipidation (Figure 4D), confirming autophagy induction to be dependent on MakA-induced aggregate formation. Furthermore, treatment of autophagy-deficient (Atg5−/−) mouse embryonic fibroblasts (MEFs) with MakA did not induce LC3 lipidation (Figure 4E), yet A568-MakA still assembled into perinuclear aggregates (Figure 4F), confirming that autophagy is not required for aggregate formation. Together, these data demonstrate that the autophagy induction observed after MakA treatment is a result of aggregate formation.

To further characterize the nature of autophagy induction MakA-induced LC3 lipidation was assessed in MEFs deficient for the ULK-interacting protein FIP200 (the key component of the ULK1/2-ATG13-FIP200 complex). FIP200 is essential for autophagosome initiation in canonical autophagy (Hara et al., 2008), but dispensable for many noncanonical autophagy pathways (Florey et al., 2011; Jacquin et al., 2017; Martinez et al., 2015). MakA-induced LC3 lipidation was maintained in the absence of FIP200 (Figure 4G) indicating activation of a noncanonical autophagy pathway. To explore it further, we employed HEK293 cells deficient for ATG16L1 and stably expressing ATG16L1α, ATG16L1β or an ATG16L1 C-terminal deletion mutant (ATG16L1 1-249). Individual ATG16L1 isoforms/mutants have been recently described to play distinct roles in the regulation of LC3 lipidation on different intracellular membranes. ATG16L1α, ATG16L1β and ATG16L1(1-249) can support LC3 lipidation in canonical autophagy, while ATG16L1(1-249) cannot support LC3-associated phagocytosis (LAP). Only the β isoform supports VPS34-independent LC3 lipidation on perturbed endosomes (Fletcher et al., 2018; Lystad et al., 2019). MakA-induced LC3 lipidation was blocked in ATG16-KO cells and restored in cells expressing the β isoform of ATG16L1, but not in cells expressing the α isoform or ATG16L1(1-249) (Figure 4H). These results suggested that MakA induces neither canonical autophagy nor LAP, but specifically induces LC3 lipidation on perturbed endosomal membranes (Lystad et al., 2019), in line with the endolysosomal composition of the membrane, and lack of double-membrane structures observed in TEM images (Figure 2A). Interestingly, MakA-induced membranous aggregates were found to be devoid of Galectin3 (Figure S5), a sensor of endomembrane damage implicated in the subsequent autophagic response (Chauhan et al., 2016).

### Sequestration of the ATG12-ATG5-ATG16L1 complex at MakA-induced aggregates inhibits both canonical autophagy and autophagy-related processes

ATG16L1, in complex with ATG5 and ATG12, localizes to the site of autophagosome initiation and functions as an E3-like enzyme to catalyze the conjugation of PE to LC3 (Fujita et al., 2008; Mizushima et al., 2003). Normally dispersed throughout the cytosol (Figure S6A), ATG16L1 and ATG12 were found to be enriched at MakA-induced aggregates after toxin treatment (Figure 5A). With no observable change in the total protein levels of ATG16L1 or ATG12 after toxin treatment (Figure S6B), we aimed to determine how sequestration of the E3-like complex might impact canonical and noncanonical autophagy pathways dependent on the same machinery. To assess canonical autophagy function, we explored the effect of MakA on autophagosome formation induced by mTOR inhibition. In vehicle-treated cells, ATG16L1 (Figure 5B) and ATG12 (Figure S6C) were dispersed throughout the cytosol. After torin1 treatment, both proteins localized to punctate structures marking the sites of autophagosome formation (Figure 5B and Figure S6C). In MakA-treated cells, ATG16L1/ATG12 were enriched at the aggregate prior to torin1 treatment. After treatment, we observed significantly less ATG16L1/ATG12 puncta as compared to vehicle-treated cells (Figure 5C and Figure S6D). Furthermore, puncta that did appear were predominantly associated with residual aggregates, marked by mono- and poly-ubiquitinylated conjugates (Figure 5D and Supplementary Video 2). These data suggest that initiation site formation is inhibited in MakA-treated cells. To confirm, Cos7 cells were transiently transfected with EGFP-LC3, pre-treated with MakA to induce aggregate formation, and treated with torin1 to induce autophagy. Cells were imaged every 30 s for 1 h to observe the total number and location of EGFP-LC3 puncta. MakA treatment not only resulted in a significant decrease in the number of EGFP-LC3 puncta, but the puncta which were quantified were predominantly associated with the existing aggregate (Fig. 5E, Supplementary Video 3 and Figure S7). These results confirm that the presence of MakA-induced endolysosomal aggregates leads to the inhibition of canonical autophagy.

**Figure 5.**
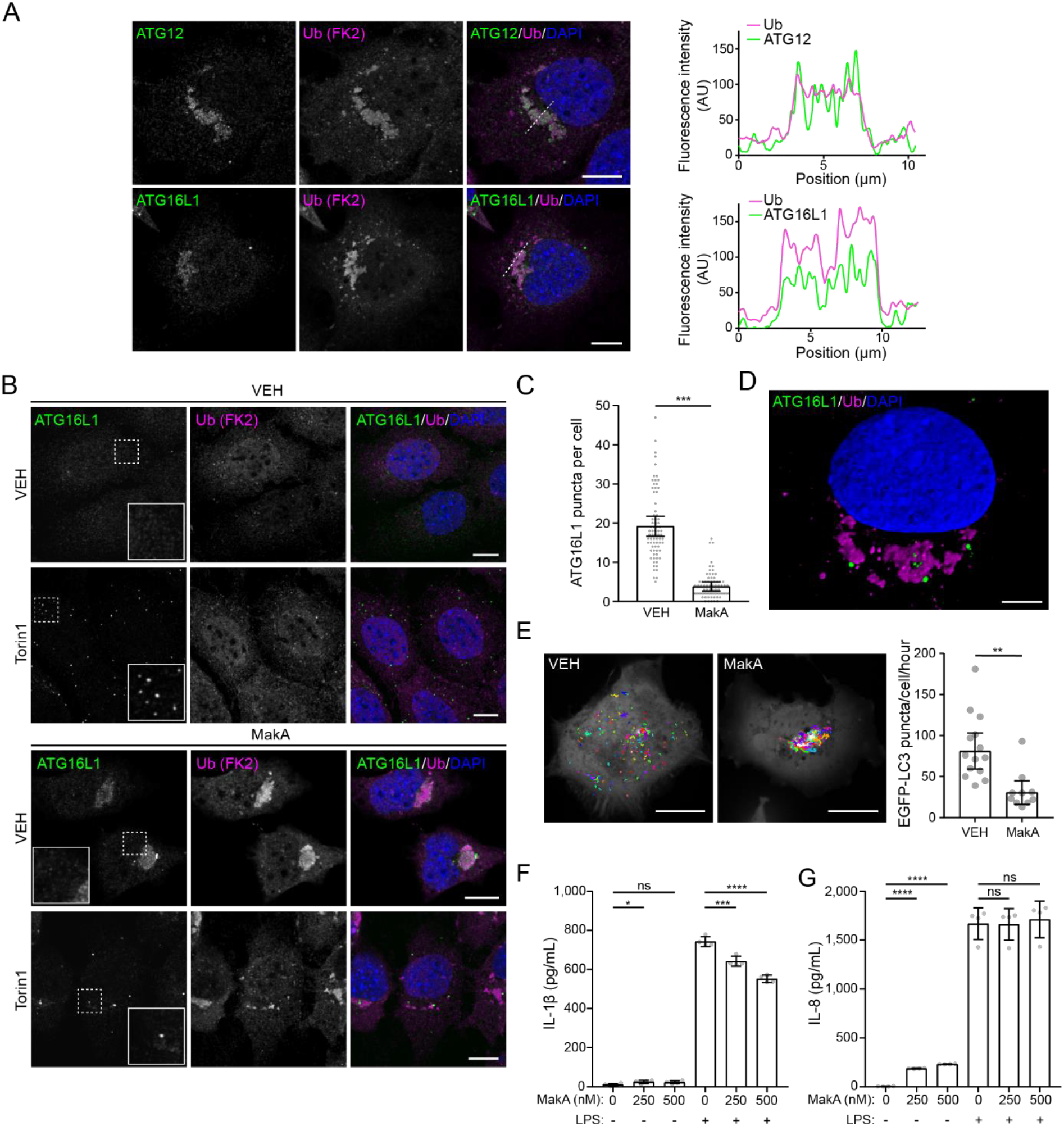
Sequestration of autophagic machinery at MakA-induced aggregates inhibits both canonical autophagy and noncanonical autophagy-related processes. (**A**) MEFs treated with 250 nM MakA for 24 h. Immunofluorescence was performed using antibodies against ATG12 (top) or ATG16L1 (bottom) and mono- and polyubiquitinylated conjugates (Ub (FK2)). Nuclei were counterstained with DAPI. Scale bars, 10 μm. Fluorescence intensity profiles along dotted white line are shown to the right of the corresponding image. (**B**) MEFs pretreated with vehicle or 250 nM MakA for 24 h followed by a 6 h treatment with vehicle or torin1 (1 μM) as indicated. Immunofluorescence was performed using antibodies against ATG16L1 and mono- and polyubiquitinylated conjugates (Ub (FK2)). Nuclei were counterstained with DAPI. Scale bars, 10 μm. (**C**) Quantification of the number of ATG16L1 puncta per cell in MEFs pretreated with vehicle or 250 nM MakA for 24 h followed by a 6 h treatment with torin1 (1 μM). Bars show mean ± sd from three biologically independent experiments. Data points represent individual cells pooled from the three independent experiments (N≥22 cells per experiment). Significance was determined from biological replicates using a two-tailed, unpaired t-test. ***p=0.0007. (**D**) 3D reconstruction of MakA-treated MEF after 6 h treatment with 1 μM torin1. Scale bar, 5 μm. (**E**) Cos7 cells transiently transfected with EGFP-LC3 were pretreated with vehicle or 250 nM MakA for 48 h then treated with 1 μM torin1 and imaged every 30 seconds for 1 hour. (left) Representative images showing the location of EGFP-LC3 puncta formation. Scale bars, 20 μM. Additional cells shown in Figure S5. (right) Total number of EGFP-LC3 puncta appearing within the hour were quantified. Bars show mean ± sd from five (VEH) or four (MakA) biologically independent experiments. Data points represent individual cells pooled from the independent experiments. Significance was determined from biological replicates using a two-tailed, unpaired t-test. **p=0.0056. (**F**, **G**) Concentration of IL-1β (F) and IL-8 (G) secreted from THP-1 cells pretreated with the indicated concentration of MakA and stimulated with 1 μg/mL LPS. Bars show mean ± sd from four biologically independent experiments. Data points represent independent experiments. Significance was determined from biological replicates using a one-way ANOVA with Sidak’s multiple comparisons tests. *p=0.0462, ***p=0.0004, ****p<0.0001, ns = not significant.

Beyond their well-established roles in regulating intracellular degradation, autophagy-related proteins (ATGs) have been implicated as regulators of a wide variety of non-autophagic cellular processes (Cadwell and Debnath, 2018). One such process with relevance to bacterial infection is the control of protein secretion (secretory autophagy) (Ponpuak et al., 2015). Secretory autophagy involves the transport of proteins via the autophagosome to the plasma membrane for extracellular release. Amongst the proteins whose release is known to be regulated by the autophagic machinery is the pro-inflammatory cytokine IL-1β (Dupont et al., 2011; Kimura et al., 2017; Zhang et al., 2015; Öhman et al., 2014); a known mediator of the innate immune response to *V. cholerae* infection (Haines et al., 2005; Kayagaki et al., 2011; Queen et al., 2015; Toma et al., 2010). To determine whether sequestration of the autophagic machinery by MakA-induced aggregates would alter IL-1β secretion, human monocytes (THP-1) were pre-treated with increasing concentrations of toxin and stimulated with lipopolysaccharide (LPS). Unlike the *V. cholerae* accessory toxins hemolysin and MARTX (Toma et al., 2010), MakA treatment was limited in its ability to induce IL-1β secretion, but did result in a significant dose-dependent inhibition of the IL-1β secretion in response to LPS stimulation (Figure 5F). Importantly, MakA did not inhibit LPS-induced secretion of IL-8 (Figure 5G) which is secreted through the autophagy-independent canonical ER-Golgi pathway and is insensitive to autophagy inhibition (Iula et al., 2018). This confirms that sequestration of the autophagic machinery at MakA-induced aggregates is capable of inhibiting canonical autophagy and autophagy-mediated processes.

## Discussion

Autophagy plays a critical role in host defense against many pathogens. As a result, evolution of pathogens has led to diverse mechanisms for evasion and subversion of host autophagy. In this study, we determine the molecular basis of the interaction between the recently identified *V. cholerae* toxin MakA and host cells, revealing a novel mechanism of autophagy subversion. After endocytosis into the host cell, MakA promotes the formation of an endolysosomal membrane-rich aggregate which becomes the target of the noncanonical autophagy pathway mediating LC3 lipidation onto perturbed endosomal membranes.

Ubiquitination and p62 recruitment were observed at the membranous aggregate yet the aggregate was found to be devoid of Galectin3, a marker of endomembrane damage, suggesting MakA-induced LC3 lipidation of endosomal membranes occurs through a Galectin3-independent mechanism (Chauhan et al., 2016). We found that MakA specifically induces the noncanonical autophagy pathway (not LAP), which was shown to be dependent on the membrane binding region and the WD-repeat containing C-terminal domain of ATG16L1 (Fletcher et al., 2018; Lystad et al., 2019). Sequestration of the E3-like ATG12-ATG5-ATG16L1 complex at the membranous aggregate resulted in the inhibition of canonical autophagy as well as autophagy-mediated cytokine secretion. How the E3-like complex is recruited to the aggregate needs further investigation but may be ubiquitin-dependent as ubiquitin has been shown previously to bind the WD-repeat domain of ATG16L1 to enhance the recruitment of ATG16L1 to damaged endosomes during bacterial infection (Fujita et al., 2013).

As compared to multiple intracellular pathogens that are targeted by and subvert xenophagy, it is less clear why a noninvasive bacterium such as *V. cholerae* has evolved to regulate autophagy (McEwan, 2017; Wu and Li, 2019). Nonetheless, its importance is emphasized by the fact that several *V. cholerae* toxins have evolved to engage host cell autophagy. The pore-forming exotoxin *V. cholerae* cytolysin (VCC) has been shown to induce autophagy (Elluri et al., 2014; Gutierrez et al., 2007), the α/β-hydrolase effector domain of multifunctional-autoprocessing repeats-in-toxin (MARTX) toxin has PI3P-specific phospholipase A1 (PLA1) activity capable of inhibiting autophagosome formation by reducing intracellular PI3P levels (Agarwal et al., 2015), and cholera toxin has been shown to inhibit autophagy through increasing cyclic AMP production (Shahnazari et al., 2011). MakA is unique in that it induces noncanonical host-cell autophagy resulting in a spatial inhibition of canonical autophagy with autophagosome formation restricted to within close proximity of the toxin-induced perinuclear aggregate. How, or if, these different autophagy-modulating toxins of *V. cholerae* may work in concert remains unknown, but our data suggest that the combined disruption of autophagy could serve to dampen the innate immune response to bacterial infection. The MakA protein is a factor found in all pathotypes of *V. cholerae* and it may be of particular relevance for the pathogenesis of *V. cholerae* that are causing not only the diarrheal disease cholera, but are responsible for severe extra-intestinal infections and bacteremia.

The narrow pH window in which MakA is shown to induce aggregate formation could provide site-specificity within the human gastrointestinal tract due a natural pH gradient between the stomach and colon (Cook et al., 2012). The observed pH range at which MakA induced aggregate formation *in vitro* closely matches that of the small intestine (pH 6.15 - 7.88), the preferential site of colonization for *V. cholerae*. Thus, we hypothesize that MakA could contribute to *V. cholerae* pathogenicity through localized innate immune suppression at the colonization site. This pH dependence also raises an important caveat to modeling *V. cholerae* infection in mice as mice have a lower intestinal pH than humans (McConnell et al., 2008). With a mean intestinal pH below 5, bacterial toxins that have evolved to function within a pH range relevant to the human intestinal tract would be inactive in mice. In agreement with this hypothesis, a Δ*makA V. cholerae* mutant with reduced pathogenicity in *C. elegans* and zebrafish (Dongre et al., 2018) was not limited in its ability to colonize mice, nor did it impact cytokine secretion (Figure S78). Thus, care must be taken when translating *V. cholerae* infection data from mouse to humans.

Juxtanuclear aggregate formation has been reported in the cellular response to various pathogens including *Clostridium difficile* (Miura et al., 2011), *Salmonella enterica* (López-Montero et al., 2016; Mesquita et al., 2012), prions (Kristiansen et al., 2005), and large cytoplasmic DNA viruses (e.g., iridovirus, vaccinia virus, African swine fever virus) which assemble virus factories for replication (Wileman, 2006). While the composition and assembly of each aggregate differ slightly, many have been reported to engage host autophagy (Boellaard et al., 1991; Heiseke et al., 2010; López-Montero et al., 2016; Wileman, 2006). MakA-induced endomembrane-rich aggregates most closely resemble the lysosomal membrane glycoprotein-positive (LGP^+^) aggregates induced in response to *Salmonella* infection of fibroblasts (López-Montero et al., 2016). Similarly to MakA-induced aggregates, LGP^+^ aggregates are endomembrane-rich and, although reportedly devoid of ubiquitin, engage the autophagic machinery for degradation. The aggregates simultaneously degrade intraphagosomal bacteria within their vicinity while those bacteria that do not come within close proximity of the aggregate are spared from autophagic degradation (López-Montero et al., 2016). This further supports our hypothesis that autophagic targeting of cellular aggregates can result in a spatial restriction of autophagy. It also suggests that the cytoplasmic aggregate generation observed in response to multiple pathogens may represent a common mechanism of subverting host autophagy.

## Materials and Methods

### Contact for Reagent and Resource Sharing

Further information and requests for reagents may be directed to, and will be fulfilled by, the corresponding authors Sun Nyunt Wai and Yaowen Wu.

### Experimental Model and Subject Details

#### Bacterial strains and culture conditions

The wild-type *Vibrio cholerae* O1 El Tor strain A1552 (Yildiz and Schoolnik, 1998) and A1552 Δ*makA* (Dongre et al., 2018) were used in the current study. Bacterial supernatants were isolated under non-cholera toxin producing conditions. Briefly, bacteria were grown to an optical density at 600 nm (OD_600nm_) of 2.0 in Luria-Bertani (LB) broth. One-millilitre bacterial culture was taken and centrifuged at 8,000 x *g* for 5 min. The supernatant was isolated and filtered through a 0.45-μm polyvinylidene difluoride (PVDF) membrane filter (Millipore, Merck Chemicals and Life Science, Solna, Sweden). The resulting culture supernatant after filtration was used for the treatment of HCT8 cells.

#### Mammalian cell lines and culture conditions

HCT8 (XY) (ATCC), THP-1 (XY) (ATCC) and mouse embryonic fibroblasts (MEFs) (Atg5 −/−: kind gift of Noboru Mizushima –Tokyo Medical and Dental University, Tokyo, Japan) (Kuma et al., 2004) (FIP200 −/−: kind gift of Jun-Lin Guan, University of Cincinnati, USA) (Gan et al., 2006) were cultured in Roswell Park Memorial Institute (RPMI) 1640 medium (Sigma-Aldrich) supplemented with 10% fetal bovine serum (FBS), 1% penicillin/streptomycin, and non-essential amino acids at 37°C with 5% CO_2_. Cos7 (ATCC), HeLa (ATCC), HEK293 ATG16-KO (Lystad et al., 2019) and HEK293T (ATCC) cells were cultured in Dulbecco’s modified Eagle medium (DMEM) (Sigma-Aldrich) supplemented with 10% fetal bovine serum (FBS), 1% penicillin/streptomycin, and non-essential amino acids at 37°C with 5% CO_2_. Cells were routinely tested for mycoplasma contamination using the LookOut mycoplasma PCR detection kit (Sigma-Aldrich). Cell lines were authenticated by ATCC with no further authentication carried out after purchase.

#### Mouse colonization

Animal experiments were performed in accordance with the animal protocols that were approved by the Ethical Committee of Animal Experiments of Nanjing Agricultural University (permit SYXK [Su] 2017-0007).

The streptomycin-treated adult mouse model was used as previously described (Wang et al., 2017). Briefly, five-week-old CD-1 mice were provided with drinking water containing 0.5% (wt/vol) streptomycin and 0.5% aspartame for 12 h before inoculation. For single colonization, approximately 10^8^ cells of wild type (SmR, lacZ-) or Δ*makA* mutant (SmR, lacZ+) mutant were intragastrical inoculated into each mouse. Fecal pellets were collected at the indicated time points, resuspended in LB broth, serially diluted and spread on plates containing X-Gal. Colony-forming unit (CFU) was counted after 24 h.

### Method Details

#### Antibodies

EEA1 (#3288, IF: 1:100), Rab7 (#9367, IF: 1:100), Rab11 (#5589, IF: 1:100), LC3B (#2775, WB: 1:1000, IF: 1:200), LAMP1 (#9091, IF: 1:200), ATG16L1 (#8089, IF: 1:100), GM130 (#12480, IF: 1:200), ATG12 (mouse specific) (#2011, WB: 1:1000, IF: 1:100), ATG12 (human specific) (#2010, WB: 1:1000, IF: 1:100), Cav1 (#3267, IF: 1:400), Galectin-3 (#87985, IF: 1:400) and Na/K-ATPase α1 (#23565, IF: 1:100) antibodies were purchased from Cell Signaling Technology. Anti-beta-actin antibody (A2228, WB: 1:10,000) was purchased from Sigma-Aldrich. Anti-p62/SQSTM1 (PM045, WB: 1:1000, IF: 1:200) was purchased from MBL international. Mono- and polyubiquitinylated conjugates (FK2) antibody (BML-PW8810, IF: 1:100) was purchased from Enzo Life Sciences. Goat anti-rabbit-HRP (Cat# 31460, WB: 1:10,000) and goat anti-mouse-HRP (Cat# 31430, WB: 1:10,000) antibodies were purchased from Thermo Fisher. Alexa Fluor 488/594/647 conjugated secondary antibodies for immunofluorescence were purchased from Jasckson ImmunoResearch.

#### Plasmids and transfection

EGFP-Rab5 construct was generated by PCR amplifying human Rab5a from cDNA and subcloning into the pEGFP-C2 (Clontech) vector using XhoI/BamHI restriction sites. EGFP-Rab5 S34N construct was generated via site-directed mutagenesis. The EGFP-LC3 plasmid has been described previously (Laraia et al., 2019). Transfection of DNA constructs was performed using X-tremeGENE HP transfection reagent (Roche) according to the manufacturer’s directions.

#### MakA purification and labeling

MakA expression and purification was carried out as previously described (Dongre et al., 2018). Alexa Fluor 568 labeling of MakA was performed using an Alexa Fluor 568 protein labeling kit (Thermo Fisher) according to the manufacturer’s instruction.

#### Cell viability assay

Cell viability of HCT8 cells treated with MakA for 48 h was quantified by PrestoBlue (Thermo Fisher) or MTS (Promega) cell viability assays, according to manufacturers instruction. PrestoBlue fluorescence (Ex/Em = 560/590 nm) and MTS absorbance (490 nm) were measured on an Infinite M200 microplate reader (Tecan). Data from both assays were normalized to untreated cells and expressed as a percentage of the control.

#### RNA-sequencing and bioinformatics analysis

HCT8 cells were treated with MakA (500 nM) for 48 hours and total RNA was extracted using RNeasy Mini Kit (Qiagen). Library preparation and sequencing was completed by Novogene (China) with a paired-end protocol and read length of 150 nt (PE150), resulting in a total output of 20 million per sample. All reads outputs were checked for passage of Illumina quality standards (Ewels et al., 2016; Wingett and Andrews, 2018). Raw data were cleaned up with trimmomatic/0.36 (Bolger et al., 2014) to remove sequences originating from Illumina adaptors and low quality reads. Files were aligned with the human reference genome hg38 (Rhead et al., 2010) with tophat/2.1.1 (Trapnell et al., 2009) using the default options. Once the alignment was completed, samtools/1.8 (Li et al., 2009) was used to select the reads that were aligned in proper pairs, and sorted them. The reads counting per gene was performed with htseq/0.9.1 (Anders et al., 2015) using stranded mode and exon as a feature. Differential expression analysis was performed with edgeR (Robinson et al., 2010). Genes with adjusted p-values <0.05 and log2 fold changes >2 were considered differentially expressed and were functionally characterized using the DAVID Database for Annotation, Visualization and Integrated Discovery (DAVID) (Huang et al., 2009a, b) and Gene Set Enrichment Analysis (GSEA) (Mootha et al., 2003; Subramanian et al., 2005).

#### Cholesterol quantification

The total cholesterol content of HCT8 cells treated with MakA was determined using the Amplex Red Cholesterol Assay kit (Molecular Probes) as previously described (Eskelinen et al., 2004). Cells were washed with cold PBS and lysed (50 mM Tris pH 8, 2 mM CaCl_2_, 80 mM NaCl, 1% Triton X-100). For cholesterol quantification, 25 μl of lysate was mixed 1:1 with reaction buffer and then with an equivalent volume of Amplex Red working solution (300 μM Amplex Red reagent, 2 U/mL HRP, 2 U/mL cholesterol oxidase, 0.2 U/mL cholesterol esterase). Reactions were incubated for 30 minutes at 37 °C and fluorescence measured using a Synergy H4 microplate reader (BioTek) (Ex/Em = 535/595 nm). Cholesterol content was normalized to total protein, as determined by Bradford assay (Bio-Rad Protein Reagent Dye).

#### Cholesterol staining

Free cholesterol was visualized using fillipin (Sigma-Aldrich). For staining, cells were fixed in 3% paraformaldehyde in phosphate-buffered saline (PBS) for 1 h at room temperature, washed 3 times with PBS containing 1.5 mg/mL glycine, and incubated with 0.05 mg/mL filipin in PBS containing 10% FBS for 30 minutes at 37 °C. When combined with antibody staining, cells were incubated with primary antibody immediately after filipin staining without any additional permeabilization. Primary antibody was diluted in 0.05 mg/mL filipin in PBS containing 10% FBS.

#### Correlative light and electron microscopy

Cos7 cells were plated on gridded 35 mm Matek dishes (MatTek Corporation P35G-1.5–14-CGRD), treated with 250 nM A568-MakA for 48 h in DMEM medium and initially fixed with paraformaldehyde (4%) and glutaraldehyde (0.5%) for 1 h at room temperature. For confocal microscopy, the cells were imaged on a Leica SP8 inverted confocal system (Leica Microsystems) equipped with a HC PL APO 63x/1.40 oil immersion lens.

After confocal microscopy, the cells were fixed in 0.05% malachite green oxalate, 2.5% glutaraldehyde and 0.1 M sodium cacodylate buffer and post-fixed with 0.8 % K3FC(CN)_6_ and 1% OsO4 (14 min). Samples were further stained with 1% tannic acid and 1% uranyl acetate (Polysciences Inc., Hirschberg an der Bergstrasse, DE) and then left in 100% resin for 1 hr at room temperature and later polymerised overnight at 60°C. After polymerization blocks were trimmed down to the regions previously identified on the confocal microscope based on the grid markings imprinted on the resin. Sections were cut on an Ultracut UCT ultramicrotome (Leica) and collected on formvar-coated slot grids. Samples were observed in Thermo Scientific™ Talos L120C transmission electron microscope (TEM). Image overlay of immunofluorescence images and electron micrographs was performed manually using Adobe Photoshop.

#### Immunofluorescence

For fixed cell immunofluorescence, cells were grown on poly-L-lysine coated coverslips and fixed either in 3% paraformaldehyde in PBS for 10 min at room temperature or with ice-cold methanol for 15 min at −20 °C. Cells were then washed 3 times with PBS containing 1.5 mg/mL glycine, permeabilized in 0.25% Triton X-100 in PBS for 5 min and washed 3 times with PBS (Note: permeabilization step was skipped with methanol fixation or when immunofluorescence was preceded by filipin staining). Cells were blocked with 5% donkey serum for 30 min followed by a 1-2 h incubation with primary antibody at room temperature. Cells were then washed 3 times with PBS and incubated with Alexa Fluor conjugated secondary antibodies for 30 min at room temperature. Cells were washed 3 times with PBS and mounted on slides using ProLong Diamond antifade mountant (Thermo Fisher). For live-cell imaging, cells were seeded on poly-L-lysine coated 8-well cover glass-bottom chamber slides (Sarstedt) and incubated for 24 h. Imaging was performed in DMEM/RPMI without phenol red (Sigma-Aldrich).

Immunofluorescence imaging was performed on two microscope systems: 1) Zeiss Cell Observer spinning disk confocal (ANDOR iXon Ultra) (Carl Zeiss) equipped with a 63x immersion oil objective lens (Plan-Apochromat 1.40 Oil DIC M27) and a temperature-controlled hood maintained at 37°C and 5% CO_2_. 2) Leica SP8 FALCON inverted confocal system (Leica Microsystems) equipped with a HC PL APO 63x/1.40 oil immersion lens and a temperature-controlled hood maintained at 37°C and 5% CO_2_. 4′,6-diamidino-2-phenylindole (DAPI) and filipin were excited using a 405 nm Diode laser, and EGFP/Alexa488, mCherry/Alexa594 and Alexa-647 fluorescence were excited using a tuned white light laser. Scanning was performed in line-by-line sequential mode.

Image processing was restricted to brightness/contrast adjustment using ImageJ – FIJI distribution (Schindelin et al., 2012) (NIH). Fluorescence intensity profiles were generated using the plot profile command in ImageJ. Live-cell particle tracking was performed using the spot tracking function (Chenouard et al., 2013) of the open-source bioimage processing software, Icy (de Chaumont et al., 2012).

#### Immunoblotting

Cells were rinsed with PBS, lysed in ice-cold lysis buffer (20 mM Tris-HCl pH 8, 300 mM KCl, 10% Glycerol, 0.25% Nonidet P-40, 0.5 mM EDTA, 0.5 mM EGTA, 1 mM PMSF, 1x complete protease inhibitor (Roche)), passed 6X through a 21G needle, and cleared by centrifugation (25 min/12,700 rpm/4°C). Protein concentrations were determined via Bradford assay (Bio-Rad Protein Reagent) and lysates normalized. Lysates were then mixed with 4x sample buffer and boiled for 10 min prior to separation by SDS-PAGE and transferred to a nitrocellulose membrane (Bio-Rad) using a Trans-Blot Turbo transfer system (Bio-Rad). After a 1 h block in TBST with 5% skim milk (at room temperature), membranes were incubated with primary antibody overnight at 4°C. Membranes were then washed with TBST and incubated for 1 h at room temperature with the appropriate HRP-conjugated secondary antibody in blocking buffer. Protein detection was carried out using chemiluminescence (Bio-Rad) and imaged using a ChemiDoc imaging system (Bio-Rad).

#### Cytokine quantification

THP-1 cells were treated with MakA in a dose dependent manner for 12 h and challenged with LPS (1 μg/mL) from *E. coli* 0111:B4 (Sigma-Aldrich) for 4 h. Supernatants collected after treatment were filtered with 0.2μm Minsart® sterile syringe filter (Sartorius stedim biotech) and cytokines IL-8 (Abcam, ab214030) and IL-1ß (Abcam, ab214025) were measured via ELISA according to manufactures instruction. For mouse IL-1ß measuremements, heart blood was collected at 24 hrs post-infection by cardiac puncture and allowed to clot overnight at 4°C. Serum was collected by centrifugation at 3000 rpm for 15 min at 4 °C and stored at −20°C. IL-1ß was measured with an IL-1ß ELISA kit (Jiangsu Meimian Industrial Co., Ltd, China). Four independent experiments were performed.

### Quantification and statistical analysis

Data are shown as mean ± s.d. Statistical significance was determined by one/two-way ANOVA or by Student’s t tests (two-tailed, unpaired), as indicated in the corresponding figure legend, using GraphPad Prism v.7. *p < 0.05, **p < 0.01, ***p < 0.001, ns = not significant. No statistical methods were used to predetermine sample size. A standard sample size of n ≥ 3 was chosen to properly observe variance and ensure reproducibility. At least 3 biological replicates were carried out for all experiments, where possible. All attempts at replication were successful with no data excluded.

### Data and code availability

The RNA-Seq data produced in this study is available from the Gene Expression Omnibus (GEO), accession no. GSEXXXXXX.

## Acknowledgements

This work was supported by the European Research Council (ChemBioAP), Vetenskapsrådet (Nr. 2018-04585), and The Knut and Alice Wallenberg Foundation to Y.W.W. and by Vetenskapsrådet (VR-MH) (Nr. 2018-02914), Cancerfonden (2017-419), and Kempestiftelsernas (JCK-1728) to S.N.W‥ D.P.C. was supported by a fellowship from the Canadian Institute of Health Research (MFE-152550). K.P. acknowledges funding from Vetenskapsrådet (VR-NT) (Nr. 2016-05009). A.P. acknowledges funding from Kempe Foundation (JCK-1528) and the Knut and Alice Wallenberg Foundation (KAW 2015.0225). A.S. and A.H.L. were supported by the Research Council of Norway through its Centres of Excellence funding scheme (Project: 262652). We acknowledge Noboru Mizushima for sharing the Atg5 −/− MEFs and Jun-Lin Guan for sharing the FIP200 −/− MEFs. We thank Nicholas Ktistakis for help on distribution of the FIP200 −/− MEFs. The computations were performed on resources provided by SNIC through Uppsala Multidisciplinary Center for Advanced Computational Science (UPPMAX) under Project SNIC 2017-7-258. We acknowledge the Protein Expertise Platform (PEP) at Umeå University for construct design and cloning. We acknowledge the Biochemical Imaging Center (BICU) at Umeå University and the National Microscopy Infrastructure, NMI (VR-RFI 2016-00968) for providing assistance in microscopy.

## Author contributions

D.P.C. carried out cholesterol quantification/visualization, endocytosis studies, autophagy studies, and wrote the initial manuscript with input from all authors. A.N. carried out RNA sequencing sample prep and data interpretation, initial experiment on LC3 lipidation, CLEM, cytokine quantification and cell viability assays. A.H. and K.M.A. assisted with western blotting. R.C.R and A.P. analysed RNA sequencing data. K.P. purified MakA. TL and HW performed mouse colonization assay. A.S. and A.H.L. provided the HEK293 ATG16L1KO and ATG16L1 rescue cell lines. All authors commented on the manuscript. Y.W.W., S.N.W. and B.E.U. supervised the study. D.P.C., A.N., Y.W.W and S.N.W. discussed all results and finalized the manuscript.

---

**Figure S1.**
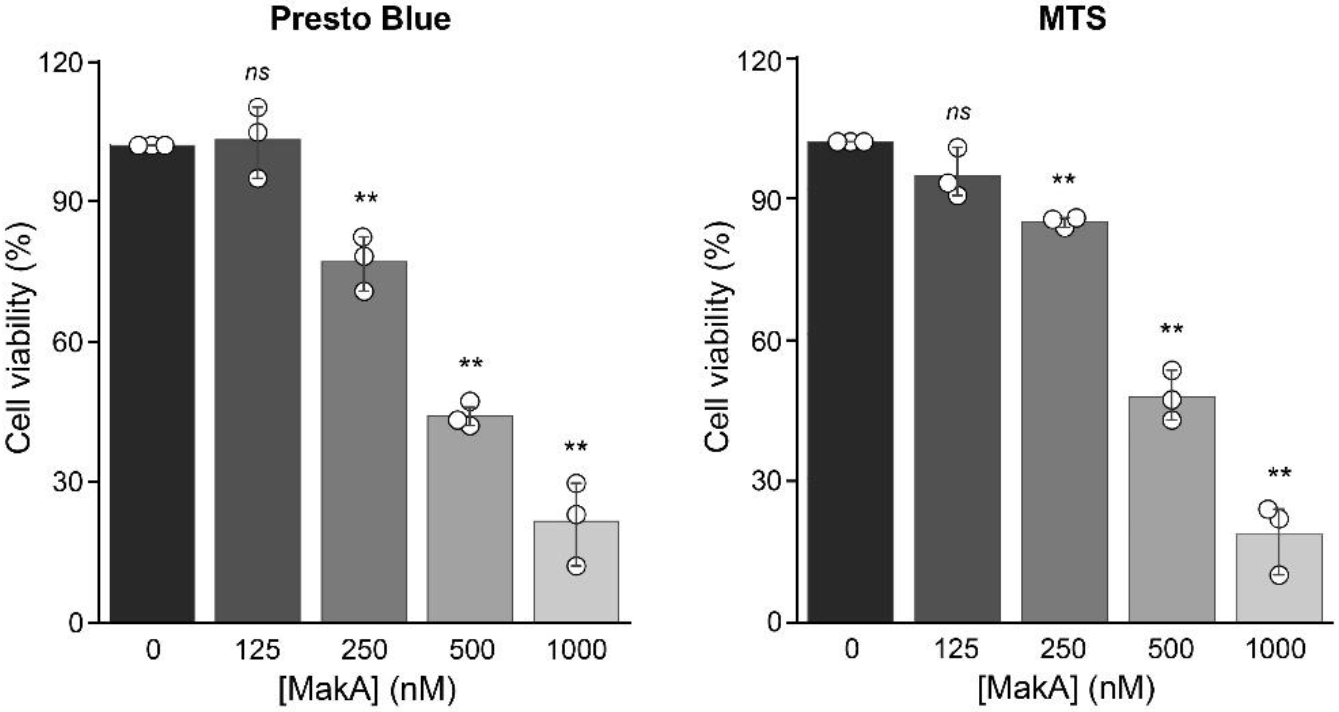
Assessment of MakA cytotoxicity in HCT8 cells. HCT8 cells treated with the indicated concentration of MakA for 48 h and toxicity assessed by Presto Blue (left) and MTS (right) cell viability assays. Data points represent three biologically independent experiments; bar graphs show mean ± s.d. Significance was determined from biological replicates using a two-tailed, unpaired t-test. ns = not significant, **p<0.01.

**Figure S2.**
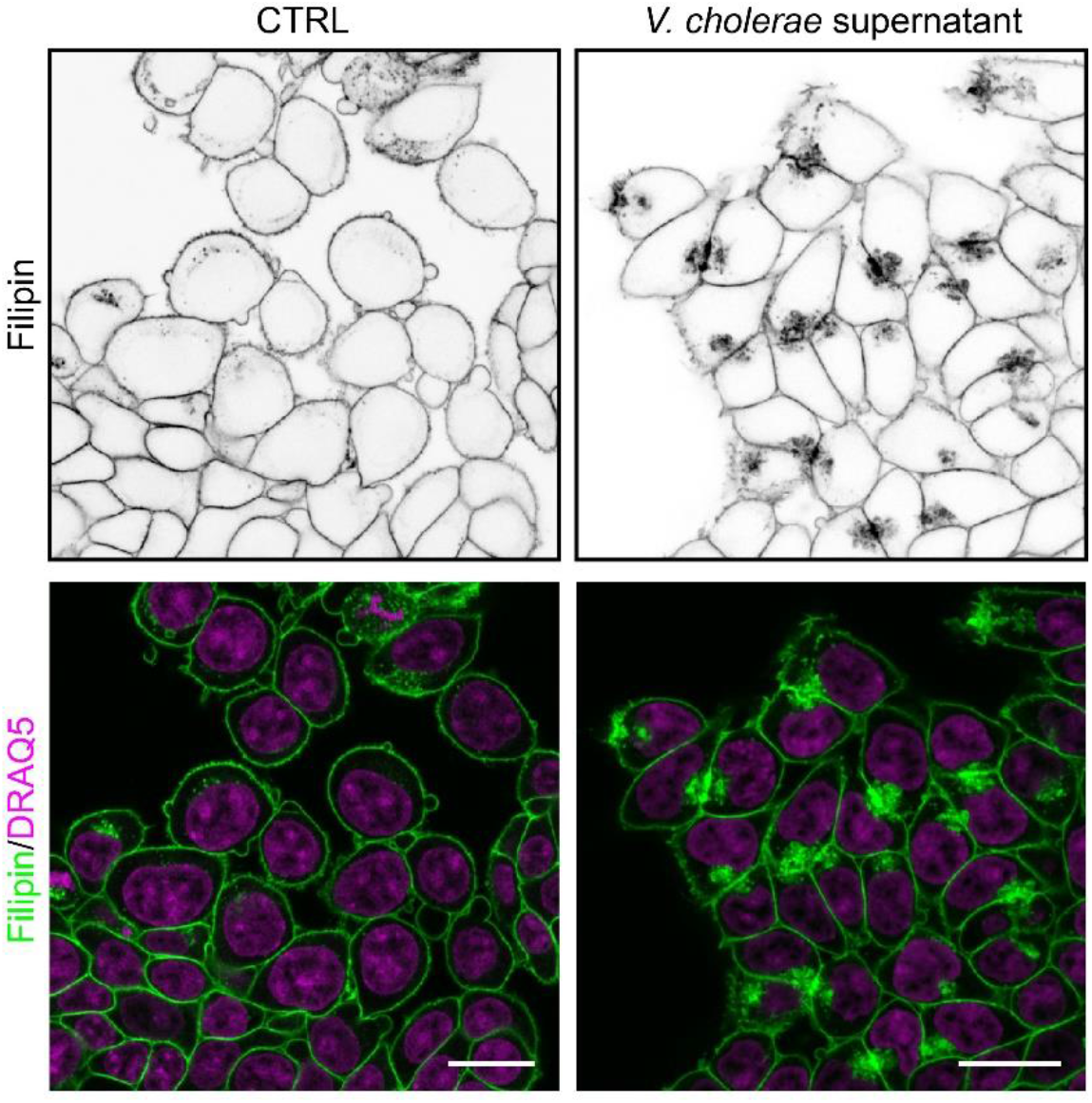
Filipin staining of HCT8 cells treated with supernatant collected from *V. cholerae* cultures. Intracellular free cholesterol localization following 6 h treatment with supernatant collected from *V. cholerae*, as detected by filipin staining. Nuclei were counterstained with DRAQ5. Scale bars, 20 μm.

**Figure S3.**
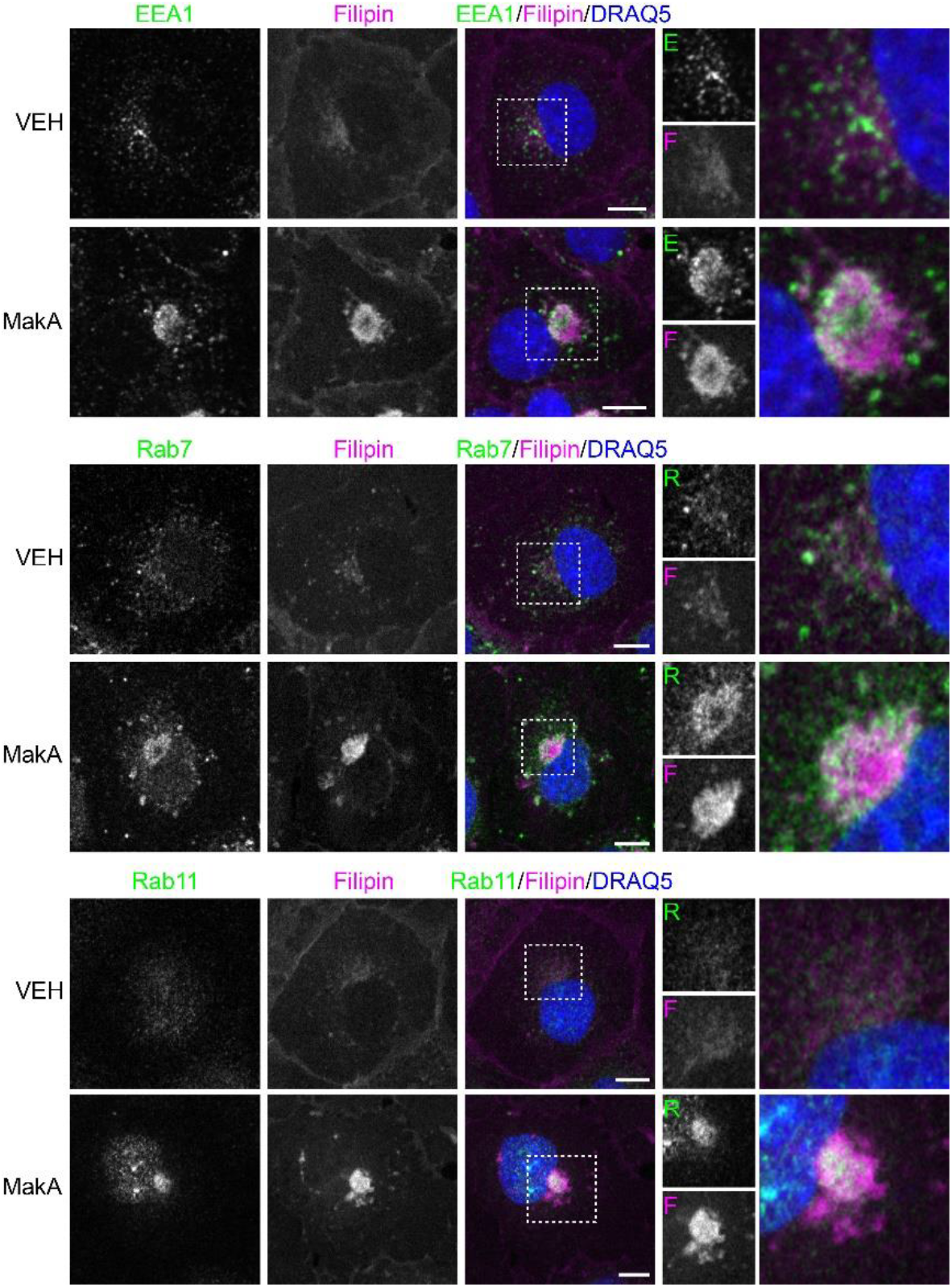
MakA induced aggregates are enriched in endolysosomal membranes. Cos7 cells treated with vehicle (VEH) or 250 nM MakA for 48 h. Cells were stained with filipin to visualize MakA-induced aggregates and immunofluorescence performed with antibodies against EEA1, Rab7 or Rab11 to visualize early, late and recycling-endosomes, respectively. Nuclei were counterstained with DRAQ5. Scale bars, 10 μm. Areas marked with dotted lines in the overlay images (a-d) are also displayed separately to show immunofluorescence detection of EEA1 (E) and Rab (R) proteins and as an enlarged image.

**Figure S4:**
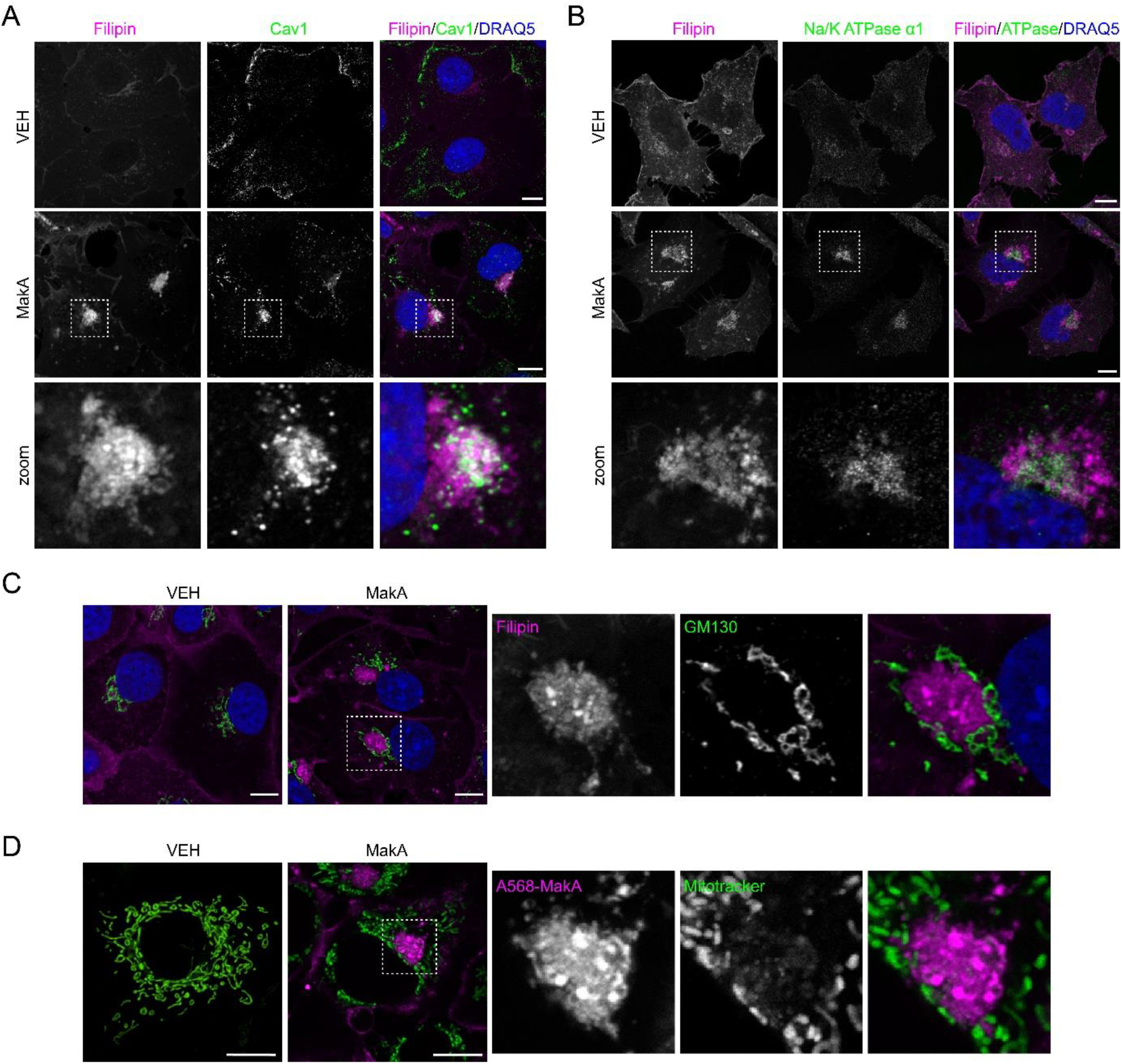
MakA induced aggregate membranes are derived from the plasma membrane. (**A, B, C**) Cos7 cells treated with vehicle (VEH) or 250 nM MakA for 48 h. Cells were stained with filipin to visualize MakA-induced aggregates and immunofluorescence performed with antibodies against Cav1 (A), Na/K ATPase α1 (B), or GM130 (C). Nuclei were counterstained with DRAQ5. Scale bars, 10 μm. (**D**) Cos7 cells treated with vehicle (VEH) or 250 nM A568-MakA for 48 h. Cells were stained with MitoTracker to visualize mitochondrial membranes. Scale bars, 10 μm.

**Figure S5:**
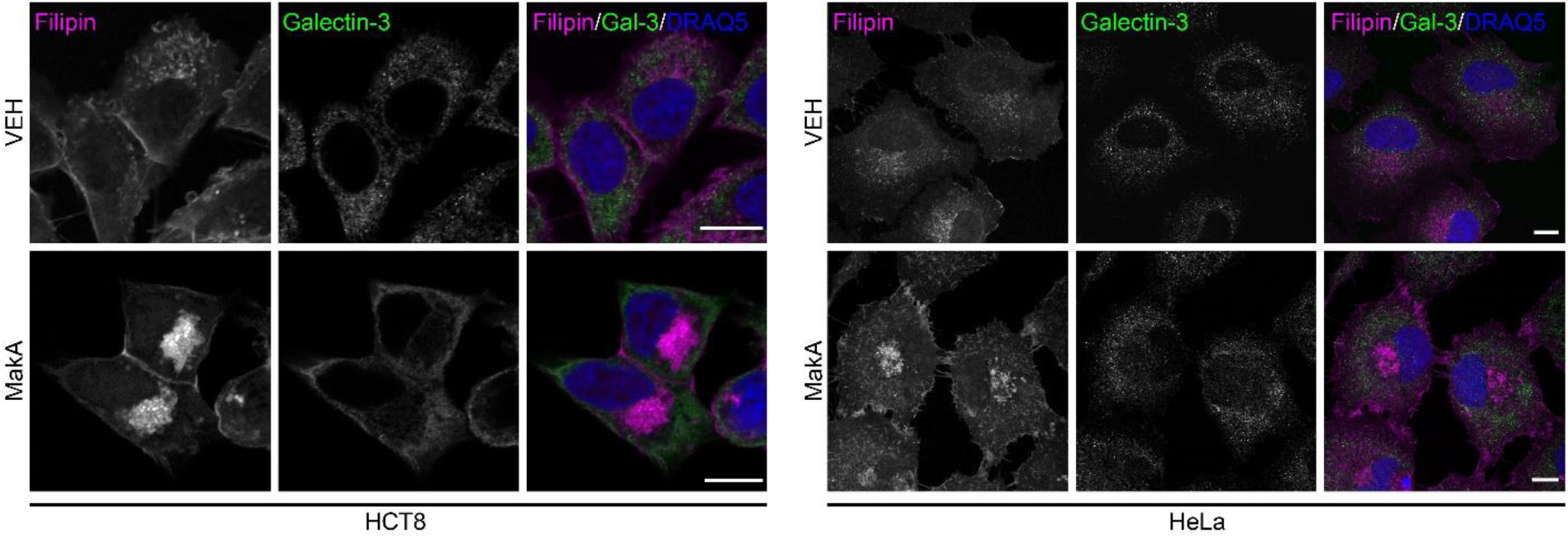
Galectin3 is not recruited to MakA induced aggregates. HCT8 (left) and HeLa (right) cells treated with 250 nM MakA for 24h (pH 7.2). Cells were stained with filipin to visualize MakA-induced aggregates and immunofluorescence performed with an antibody against Galectin3. Nuclei were counterstained with DRAQ5. Scale bars, 10 μm.

**Figure S6:**
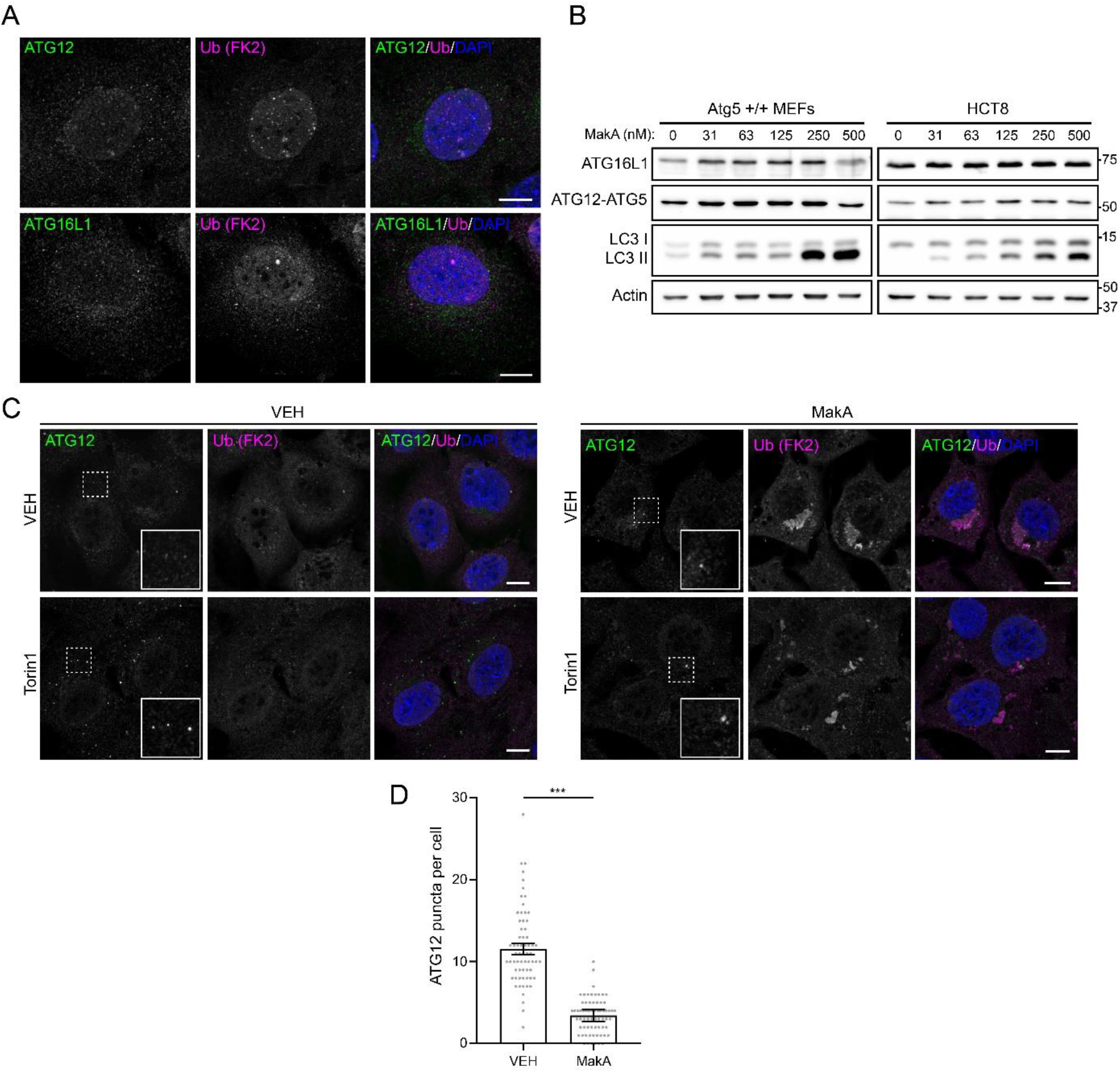
ATG12-ATG5-ATG16L1 complex response to MakA. (**A**) MEFs treated with vehicle for 24 h. Immunofluorescence was performed using antibodies against ATG12 (top) or ATG16L1 (bottom) and mono- and polyubiquitinylated conjugates (Ub (FK2)). Nuclei were counterstained with DAPI. Scale bars, 10 μm. (**B**) Western blot analysis of Atg5 +/+ MEF and HCT8 cells treated with the indicated concentration of MakA for 24 h. (**C**) MEFs pretreated with vehicle or 250 nM MakA for 24 h followed by a 6 h treatment with vehicle or torin1 (1 μM) as indicated. Immunofluorescence was performed using antibodies against ATG12 and mono- and polyubiquitinylated conjugates (Ub (FK2)). Nuclei were counterstained with DAPI. Scale bars, 10 μm. (**D**) Quantification of the number of ATG12 puncta per cell in MEFs pretreated with vehicle or 250 nM MakA for 24 h followed by a 6 h treatment with torin1 (1 μM). Bars show mean ± sd from three biologically independent experiments. Data points represent individual cells pooled from the three independent experiments (N≥20 cells per experiment). Significance was determined from biological replicates using a two-tailed, unpaired t-test. ***p=0.0001.

**Figure S7:**
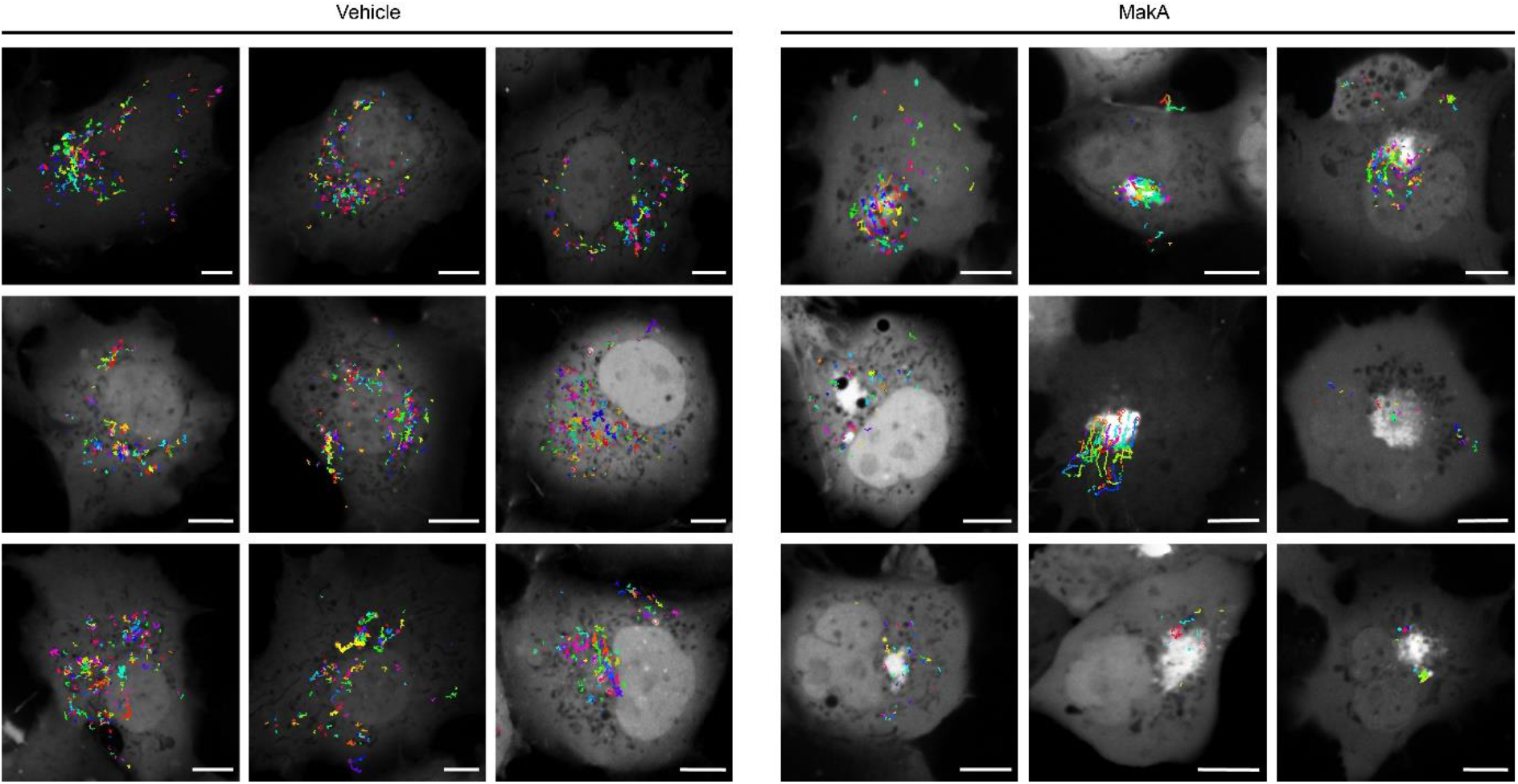
Additional Cos7 EGFP-LC3 puncta locations. Images showing the location of EGFP-LC3 puncta formation in Cos7 cells transiently transfected with EGFP-LC3, pretreated with vehicle (left) or 250 nM MakA (right) for 48 h then treated with 1 μM torin1 and imaged every 30 seconds for 1 hour. Scale bars, 10 μm.

**Figure S8:**
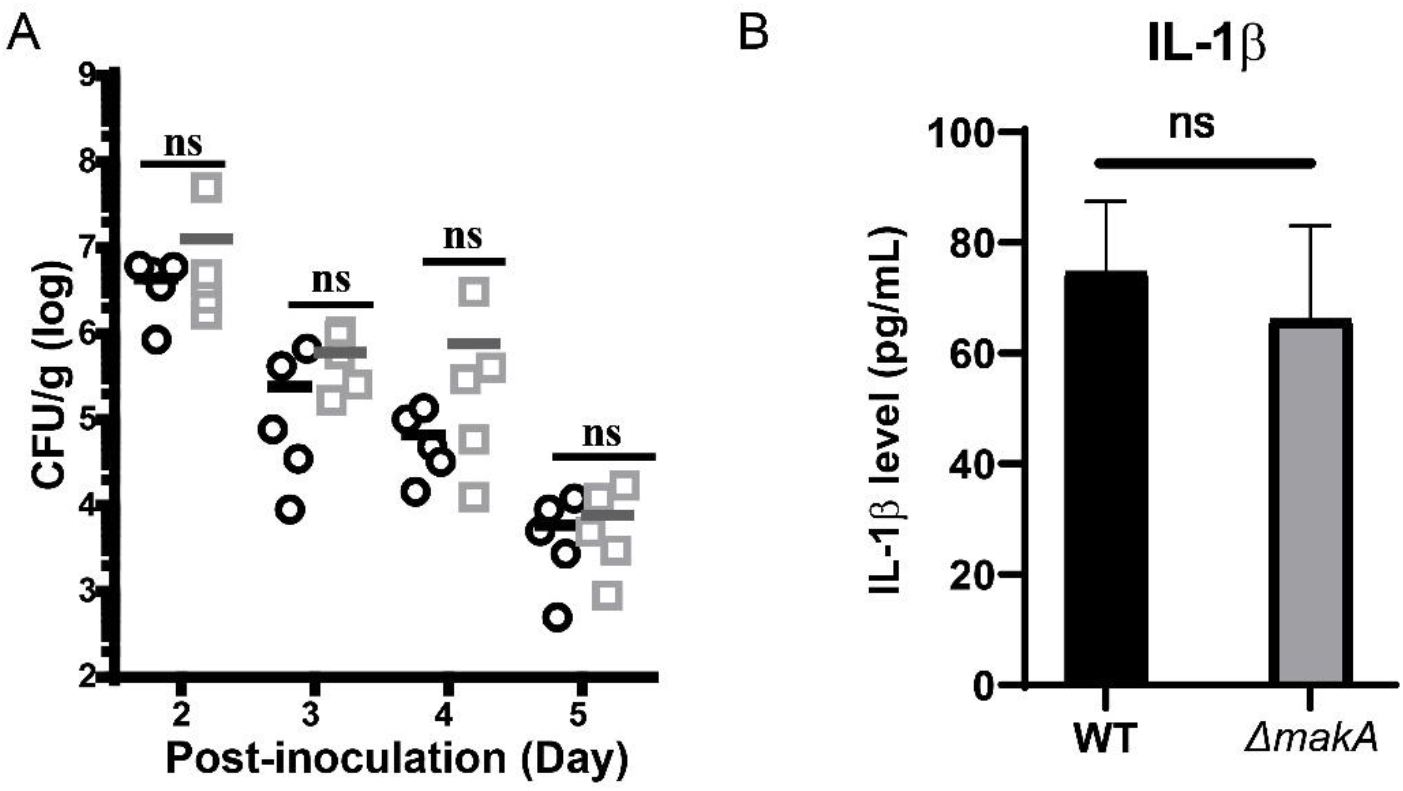
MakA does not contribute to pathogenicity in a mouse model of *V. cholerae* infection. (**A**) Five-week-old CD-1 mice were treated with streptomycin. Approximately 10^8^ cells of wild-type (lacZ-) and ΔmakA mutant (lacZ+) were respectively intragastrically administered to each mice. Fecal pellets were collected from each mouse and plated onto selective plates. CFU was counted after 24 h. Horizontal line represent means from five mice. NS, not significant. ○:WT; □: Δ*makA* mutant. (**B**) Concentration of IL-1β measured was measured in heart blood serum after 24 hrs post-infection using an ELISA kit. The data consist of the means and standard deviations from four independent experiments. *, p<0.05 (Student’s t-test). NS, not significant.

**Figure S9:**
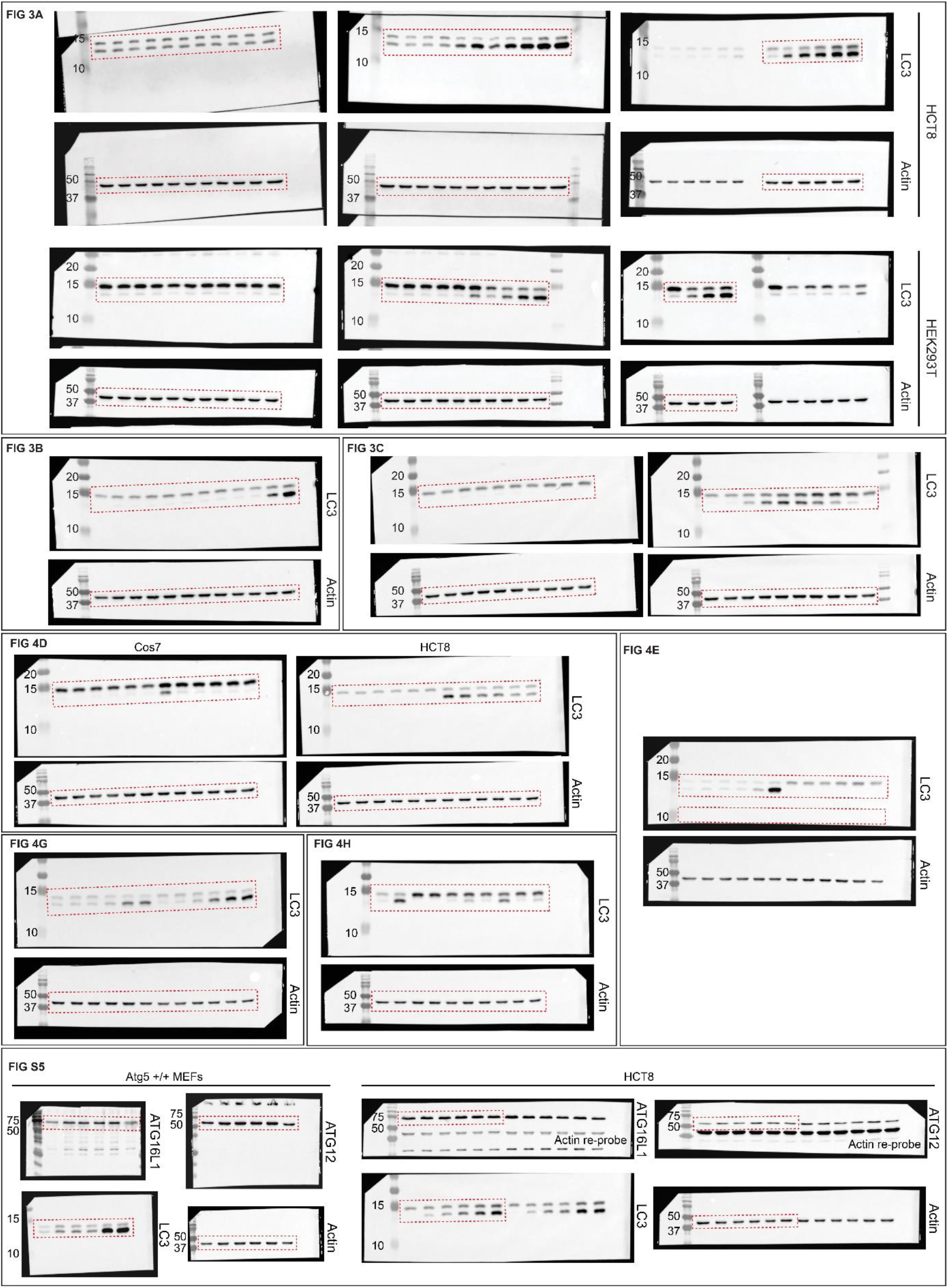
Uncropped western blots.

**Supplementary Video 1. Live-cell imaging of Cos7 cells transfected with EGFP-Rab5 and treated with 250 nM Alexa568-MakA for 48 h.** Related to Figure 2C. Frames captured every 1.3 seconds. Data are representative of two independent experiments. Scale bar = 10 μm.

**Supplementary Video 2. 3D reconstruction of MakA-treated MEF after 6 h treatment with 1 μM torin1.** Related to Figure 6D. Image reconstructed from 13 images covering a total depth of 3.88 μm.

**Supplementary Video 3. Live-cell imaging of Cos7 cells transiently transfected with EGFP-LC3.** Related to Figure 5H. Cells were pretreated with vehicle (left) or 250 nM MakA (right) for 48 h then treated with 1 μM torin1. Frames captured every 10 seconds. Data are representative of at least four independent experiments. Scale bar = 10 μm.

## REFERENCES

Agarwal, S., Kim, H., Chan, R.B., Williamson, R., Cho, W., Paolo, G.D., and Satchell, K.J. (2015). Autophagy and endosomal trafficking inhibition by Vibrio cholerae MARTX toxin phosphatidylinositol-3-phosphate-specific phospholipase A1 activity. Nat Commun 6, 8745.

Anders, S., Pyl, P.T., and Huber, W. (2015). HTSeq--a Python framework to work with high-throughput sequencing data. Bioinformatics 31, 166–169.

Boellaard, J.W., Kao, M., Schlote, W., and Diringer, H. (1991). Neuronal autophagy in experimental scrapie. Acta Neuropathol 82, 225–228.

Bolger, A.M., Lohse, M., and Usadel, B. (2014). Trimmomatic: a flexible trimmer for Illumina sequence data. Bioinformatics 30, 2114–2120.

Bucci, C., Parton, R.G., Mather, I.H., Stunnenberg, H., Simons, K., Hoflack, B., and Zerial, M. (1992). The small GTPase rab5 functions as a regulatory factor in the early endocytic pathway. Cell 70, 715–728.

Cadwell, K., and Debnath, J. (2018). Beyond self-eating: The control of nonautophagic functions and signaling pathways by autophagy-related proteins. J Cell Biol 217, 813–822.

Chauhan, S., Kumar, S., Jain, A., Ponpuak, M., Mudd, M.H., Kimura, T., Choi, S.W., Peters, R., Mandell, M., Bruun, J.A., et al. (2016). TRIMs and Galectins Globally Cooperate and TRIM16 and Galectin-3 Co-direct Autophagy in Endomembrane Damage Homeostasis. Dev Cell 39, 13–27.

Chenouard, N., Bloch, I., and Olivo-Marin, J.C. (2013). Multiple hypothesis tracking for cluttered biological image sequences. IEEE Trans Pattern Anal Mach Intell 35, 2736–3750.

Clemens, J.D., Nair, G.B., Ahmed, T., Qadri, F., and Holmgren, J. (2017). Cholera. Lancet 390, 1539–1549.

Cook, M.T., Tzortzis, G., Charalampopoulos, D., and Khutoryanskiy, V.V. (2012). Microencapsulation of probiotics for gastrointestinal delivery. J Control Release 162, 56–67.

de Chaumont, F., Dallongeville, S., Chenouard, N., Hervé, N., Pop, S., Provoost, T., Meas-Yedid, V., Pankajakshan, P., Lecomte, T., Le Montagner, Y., et al. (2012). Icy: an open bioimage informatics platform for extended reproducible research. Nat Methods 9, 690–696.

Deretic, V., Saitoh, T., and Akira, S. (2013). Autophagy in infection, inflammation and immunity. Nat Rev Immunol 13, 722–737.

Dongre, M., Singh, B., Aung, K.M., Larsson, P., Miftakhova, R., Persson, K., Askarian, F., Johannessen, M., von Hofsten, J., Persson, J.L., et al. (2018). Flagella-mediated secretion of a novel *Vibrio cholerae* cytotoxin affecting both vertebrate and invertebrate hosts. Commun Biol 1, 59.

Dupont, N., Jiang, S., Pilli, M., Ornatowski, W., Bhattacharya, D., and Deretic, V. (2011). Autophagy-based unconventional secretory pathway for extracellular delivery of IL-1β. EMBO J 30, 4701–4711.

Elluri, S., Enow, C., Vdovikova, S., Rompikuntal, P.K., Dongre, M., Carlsson, S., Pal, A., Uhlin, B.E., and Wai, S.N. (2014). Outer membrane vesicles mediate transport of biologically active Vibrio cholerae cytolysin (VCC) from V. cholerae strains. PLoS One 9, e106731.

English, L., Chemali, M., Duron, J., Rondeau, C., Laplante, A., Gingras, D., Alexander, D., Leib, D., Norbury, C., Lippé, R., et al. (2009). Autophagy enhances the presentation of endogenous viral antigens on MHC class I molecules during HSV-1 infection. Nat Immunol 10, 480–487.

Eskelinen, E.L., Schmidt, C.K., Neu, S., Willenborg, M., Fuertes, G., Salvador, N., Tanaka, Y., Lüllmann-Rauch, R., Hartmann, D., Heeren, J., et al. (2004). Disturbed cholesterol traffic but normal proteolytic function in LAMP-1/LAMP-2 double-deficient fibroblasts. Mol Biol Cell 15, 3132–3145.

Ewels, P., Magnusson, M., Lundin, S., and Käller, M. (2016). MultiQC: summarize analysis results for multiple tools and samples in a single report. Bioinformatics 32, 3047–3048.

Faruque, S.M., Albert, M.J., and Mekalanos, J.J. (1998). Epidemiology, genetics, and ecology of toxigenic Vibrio cholerae. Microbiol Mol Biol Rev 62, 1301–1314.

Feng, Y., Press, B., and Wandinger-Ness, A. (1995). Rab 7: an important regulator of late endocytic membrane traffic. J Cell Biol 131, 1435–1452.

Fletcher, K., Ulferts, R., Jacquin, E., Veith, T., Gammoh, N., Arasteh, J.M., Mayer, U., Carding, S.R., Wileman, T., Beale, R., et al. (2018). The WD40 domain of ATG16L1 is required for its non-canonical role in lipidation of LC3 at single membranes. EMBO J 37.

Florey, O., Kim, S.E., Sandoval, C.P., Haynes, C.M., and Overholtzer, M. (2011). Autophagy machinery mediates macroendocytic processing and entotic cell death by targeting single membranes. Nat Cell Biol 13, 1335–1343.

Fujita, N., Itoh, T., Omori, H., Fukuda, M., Noda, T., and Yoshimori, T. (2008). The Atg16L complex specifies the site of LC3 lipidation for membrane biogenesis in autophagy. Mol Biol Cell 19, 2092–2100.

Fujita, N., Morita, E., Itoh, T., Tanaka, A., Nakaoka, M., Osada, Y., Umemoto, T., Saitoh, T., Nakatogawa, H., Kobayashi, S., et al. (2013). Recruitment of the autophagic machinery to endosomes during infection is mediated by ubiquitin. J Cell Biol 203, 115–128.

Gan, B., Peng, X., Nagy, T., Alcaraz, A., Gu, H., and Guan, J.L. (2006). Role of FIP200 in cardiac and liver development and its regulation of TNFalpha and TSC-mTOR signaling pathways. J Cell Biol 175, 121–133.

Gutierrez, M.G., Saka, H.A., Chinen, I., Zoppino, F.C., Yoshimori, T., Bocco, J.L., and Colombo, M.I. (2007). Protective role of autophagy against Vibrio cholerae cytolysin, a pore-forming toxin from V. cholerae. Proc Natl Acad Sci U S A 104, 1829–1834.

Haines, G.K., Sayed, B.A., Rohrer, M.S., Olivier, V., and Satchell, K.J. (2005). Role of toll-like receptor 4 in the proinflammatory response to Vibrio cholerae O1 El tor strains deficient in production of cholera toxin and accessory toxins. Infect Immun 73, 6157–6164.

Hara, T., Takamura, A., Kishi, C., Iemura, S., Natsume, T., Guan, J.L., and Mizushima, N. (2008). FIP200, a ULK-interacting protein, is required for autophagosome formation in mammalian cells. J Cell Biol 181, 497–510.

Heiseke, A., Aguib, Y., and Schatzl, H.M. (2010). Autophagy, prion infection and their mutual interactions. Curr Issues Mol Biol 12, 87–97.

Huang, d.W., Sherman, B.T., and Lempicki, R.A. (2009a). Bioinformatics enrichment tools: paths toward the comprehensive functional analysis of large gene lists. Nucleic Acids Res 37, 1–13.

Huang, d.W., Sherman, B.T., and Lempicki, R.A. (2009b). Systematic and integrative analysis of large gene lists using DAVID bioinformatics resources. Nat Protoc 4, 44–57.

Iula, L., Keitelman, I.A., Sabbione, F., Fuentes, F., Guzman, M., Galletti, J.G., Gerber, P.P., Ostrowski, M., Geffner, J.R., Jancic, C.C., et al. (2018). Autophagy Mediates Interleukin-1β Secretion in Human Neutrophils. Front Immunol 9, 269.

Jacquin, E., Leclerc-Mercier, S., Judon, C., Blanchard, E., Fraitag, S., and Florey, O. (2017). Pharmacological modulators of autophagy activate a parallel noncanonical pathway driving unconventional LC3 lipidation. Autophagy 13, 854–867.

Kayagaki, N., Warming, S., Lamkanfi, M., Vande Walle, L., Louie, S., Dong, J., Newton, K., Qu, Y., Liu, J., Heldens, S., et al. (2011). Non-canonical inflammasome activation targets caspase-11. Nature 479, 117–121.

Kim, P.K., Hailey, D.W., Mullen, R.T., and Lippincott-Schwartz, J. (2008). Ubiquitin signals autophagic degradation of cytosolic proteins and peroxisomes. Proc Natl Acad Sci U S A 105, 20567–20574.

Kimura, T., Jia, J., Kumar, S., Choi, S.W., Gu, Y., Mudd, M., Dupont, N., Jiang, S., Peters, R., Farzam, F., et al. (2017). Dedicated SNAREs and specialized TRIM cargo receptors mediate secretory autophagy. EMBO J 36, 42–60.

Klionsky, D.J., Abdelmohsen, K., Abe, A., Abedin, M.J., Abeliovich, H., Acevedo Arozena, A., Adachi, H., Adams, C.M., Adams, P.D., Adeli, K., et al. (2016). Guidelines for the use and interpretation of assays for monitoring autophagy (3rd edition). Autophagy 12, 1–222.

Klumperman, J., and Raposo, G. (2014). The complex ultrastructure of the endolysosomal system. Cold Spring Harb Perspect Biol 6, a016857.

Kristiansen, M., Messenger, M.J., Klöhn, P.C., Brandner, S., Wadsworth, J.D., Collinge, J., and Tabrizi, S.J. (2005). Disease-related prion protein forms aggresomes in neuronal cells leading to caspase activation and apoptosis. J Biol Chem 280, 38851–38861.

Kuma, A., Hatano, M., Matsui, M., Yamamoto, A., Nakaya, H., Yoshimori, T., Ohsumi, Y., Tokuhisa, T., and Mizushima, N. (2004). The role of autophagy during the early neonatal starvation period. Nature 432, 1032–1036.

Laraia, L., Friese, A., Corkery, D.P., Konstantinidis, G., Erwin, N., Hofer, W., Karatas, H., Klewer, L., Brockmeyer, A., Metz, M., et al. (2019). The cholesterol transfer protein GRAMD1A regulates autophagosome biogenesis. Nat Chem Biol 15, 710–720.

Levine, B. (2005). Eating oneself and uninvited guests: autophagy-related pathways in cellular defense. Cell 120, 159–162.

Li, G., and Stahl, P.D. (1993). Structure-function relationship of the small GTPase rab5. J Biol Chem 268, 24475–24480.

Li, H., Handsaker, B., Wysoker, A., Fennell, T., Ruan, J., Homer, N., Marth, G., Abecasis, G., Durbin, R., and Subgroup, G.P.D.P. (2009). The Sequence Alignment/Map format and SAMtools. Bioinformatics 25, 2078–2079.

Luo, J., Yang, H., and Song, B.L. (2020). Mechanisms and regulation of cholesterol homeostasis. Nat Rev Mol Cell Biol 21, 225–245.

Lystad, A.H., Carlsson, S.R., de la Ballina, L.R., Kauffman, K.J., Nag, S., Yoshimori, T., Melia, T.J., and Simonsen, A. (2019). Distinct functions of ATG16L1 isoforms in membrane binding and LC3B lipidation in autophagy-related processes. Nat Cell Biol 21, 372–383.

López-Montero, N., Ramos-Marquès, E., Risco, C., and García-Del Portillo, F. (2016). Intracellular Salmonella induces aggrephagy of host endomembranes in persistent infections. Autophagy 12, 1886–1901.

Macia, E., Ehrlich, M., Massol, R., Boucrot, E., Brunner, C., and Kirchhausen, T. (2006). Dynasore, a cell-permeable inhibitor of dynamin. Dev Cell 10, 839–850.

Martinez, J., Malireddi, R.K., Lu, Q., Cunha, L.D., Pelletier, S., Gingras, S., Orchard, R., Guan, J.L., Tan, H., Peng, J., et al. (2015). Molecular characterization of LC3-associated phagocytosis reveals distinct roles for Rubicon, NOX2 and autophagy proteins. Nat Cell Biol 17, 893–906.

McConnell, E.L., Basit, A.W., and Murdan, S. (2008). Measurements of rat and mouse gastrointestinal pH, fluid and lymphoid tissue, and implications for in-vivo experiments. J Pharm Pharmacol 60, 63–70.

McEwan, D.G. (2017). Host-pathogen interactions and subversion of autophagy. Essays Biochem 61, 687–697.

Mesquita, F.S., Thomas, M., Sachse, M., Santos, A.J., Figueira, R., and Holden, D.W. (2012). The Salmonella deubiquitinase SseL inhibits selective autophagy of cytosolic aggregates. PLoS Pathog 8, e1002743.

Miura, M., Kato, H., and Matsushita, O. (2011). Identification of a novel virulence factor in Clostridium difficile that modulates toxin sensitivity of cultured epithelial cells. Infect Immun 79, 3810–3820.

Mizushima, N., Kuma, A., Kobayashi, Y., Yamamoto, A., Matsubae, M., Takao, T., Natsume, T., Ohsumi, Y., and Yoshimori, T. (2003). Mouse Apg16L, a novel WD-repeat protein, targets to the autophagic isolation membrane with the Apg12-Apg5 conjugate. J Cell Sci 116, 1679–1688.

Mootha, V.K., Lindgren, C.M., Eriksson, K.F., Subramanian, A., Sihag, S., Lehar, J., Puigserver, P., Carlsson, E., Ridderstråle, M., Laurila, E., et al. (2003). PGC-1alpha-responsive genes involved in oxidative phosphorylation are coordinately downregulated in human diabetes. Nat Genet 34, 267–273.

Pankiv, S., Clausen, T.H., Lamark, T., Brech, A., Bruun, J.A., Outzen, H., Øvervatn, A., Bjørkøy, G., and Johansen, T. (2007). p62/SQSTM1 binds directly to Atg8/LC3 to facilitate degradation of ubiquitinated protein aggregates by autophagy. J Biol Chem 282, 24131–24145.

Ponpuak, M., Mandell, M.A., Kimura, T., Chauhan, S., Cleyrat, C., and Deretic, V. (2015). Secretory autophagy. Curr Opin Cell Biol 35, 106–116.

Queen, J., Agarwal, S., Dolores, J.S., Stehlik, C., and Satchell, K.J. (2015). Mechanisms of inflammasome activation by Vibrio cholerae secreted toxins vary with strain biotype. Infect Immun 83, 2496–2506.

Rhead, B., Karolchik, D., Kuhn, R.M., Hinrichs, A.S., Zweig, A.S., Fujita, P.A., Diekhans, M., Smith, K.E., Rosenbloom, K.R., Raney, B.J., et al. (2010). The UCSC Genome Browser database: update 2010. Nucleic Acids Res 38, D613–619.

Robinson, M.D., McCarthy, D.J., and Smyth, G.K. (2010). edgeR: a Bioconductor package for differential expression analysis of digital gene expression data. Bioinformatics 26, 139–140.

Schindelin, J., Arganda-Carreras, I., Frise, E., Kaynig, V., Longair, M., Pietzsch, T., Preibisch, S., Rueden, C., Saalfeld, S., Schmid, B., et al. (2012). Fiji: an open-source platform for biological-image analysis. Nat Methods 9, 676–682.

Schmid, D., Pypaert, M., and Münz, C. (2007). Antigen-loading compartments for major histocompatibility complex class II molecules continuously receive input from autophagosomes. Immunity 26, 79–92.

Shahnazari, S., Namolovan, A., Mogridge, J., Kim, P.K., and Brumell, J.H. (2011). Bacterial toxins can inhibit host cell autophagy through cAMP generation. Autophagy 7, 957–965.

Simonsen, A., Lippé, R., Christoforidis, S., Gaullier, J.M., Brech, A., Callaghan, J., Toh, B.H., Murphy, C., Zerial, M., and Stenmark, H. (1998). EEA1 links PI(3)K function to Rab5 regulation of endosome fusion. Nature 394, 494–498.

Subramanian, A., Tamayo, P., Mootha, V.K., Mukherjee, S., Ebert, B.L., Gillette, M.A., Paulovich, A., Pomeroy, S.L., Golub, T.R., Lander, E.S., et al. (2005). Gene set enrichment analysis: a knowledge-based approach for interpreting genome-wide expression profiles. Proc Natl Acad Sci U S A 102, 15545–15550.

Tanida, I., Ueno, T., and Kominami, E. (2008). LC3 and Autophagy. Methods Mol Biol 445, 77–88.

Toma, C., Higa, N., Koizumi, Y., Nakasone, N., Ogura, Y., McCoy, A.J., Franchi, L., Uematsu, S., Sagara, J., Taniguchi, S., et al. (2010). Pathogenic Vibrio activate NLRP3 inflammasome via cytotoxins and TLR/nucleotide-binding oligomerization domain-mediated NF-kappa B signaling. J Immunol 184, 5287–5297.

Trapnell, C., Pachter, L., and Salzberg, S.L. (2009). TopHat: discovering splice junctions with RNA-Seq. Bioinformatics 25, 1105–1111.

Urbé, S., Huber, L.A., Zerial, M., Tooze, S.A., and Parton, R.G. (1993). Rab11, a small GTPase associated with both constitutive and regulated secretory pathways in PC12 cells. FEBS Lett 334, 175–182.

Wang, H., Naseer, N., Chen, Y., Zhu, A.Y., Kuai, X., Galagedera, N., Liu, Z., and Zhu, J. (2017). OxyR2 Modulates OxyR1 Activity and Vibrio cholerae Oxidative Stress Response. Infect Immun 85.

Wileman, T. (2006). Aggresomes and autophagy generate sites for virus replication. Science 312, 875–878.

Wingett, S.W., and Andrews, S. (2018). FastQ Screen: A tool for multi-genome mapping and quality control. F1000Res 7, 1338.

Wu, Y.W., and Li, F. (2019). Bacterial interaction with host autophagy. Virulence 10, 352–362.

Yildiz, F.H., and Schoolnik, G.K. (1998). Role of rpoS in stress survival and virulence of Vibrio cholerae. J Bacteriol 180, 773–784.

Zaarur, N., Meriin, A.B., Bejarano, E., Xu, X., Gabai, V.L., Cuervo, A.M., and Sherman, M.Y. (2014). Proteasome failure promotes positioning of lysosomes around the aggresome via local block of microtubule-dependent transport. Mol Cell Biol 34, 1336–1348.

Zhang, M., Kenny, S.J., Ge, L., Xu, K., and Schekman, R. (2015). Translocation of interleukin-1β into a vesicle intermediate in autophagy-mediated secretion. Elife 4.

Öhman, T., Teirilä, L., Lahesmaa-Korpinen, A.M., Cypryk, W., Veckman, V., Saijo, S., Wolff, H., Hautaniemi, S., Nyman, T.A., and Matikainen, S. (2014). Dectin-1 pathway activates robust autophagy-dependent unconventional protein secretion in human macrophages. J Immunol 192, 5952–5962.

